# Label-free Machine Learning Prediction of Chemotherapy on Tumor Spheroids using a Microfluidics Droplet Platform

**DOI:** 10.1101/2025.03.25.645032

**Authors:** Caroline Parent, Hasti Honari, Tiziana Tocci, Franck Simon, Sakina Zaidi, Audric Jan, Vivian Aubert, Olivier Delattre, Hervé Isambert, Claire Wilhelm, Jean-Louis Viovy

## Abstract

To rapidly evaluate the effects of anticancer treatments in 3D models, an integrated approach is proposed, combining a droplet-based microfluidic platform for spheroid formation and single-spheroid chemotherapy application, label-free morphological analysis, and machine learning to assess treatment response. Morphological features of spheroids, such as size and color intensity, are extracted and selected using the Multivariate Information-based Inductive Causation (MIIC) algorithm, and used to train a neural network for spheroid classification into viability classes, derived from metabolic assays performed within the same platform as a benchmark.

The model was tested on Ewing sarcoma cell line and patient-derived xenograft (PDX) cells, demonstrating robust performance across datasets. It accurately predicts spheroid viability, used to generate dose-response curves and to determine half-maximal inhibitory concentration (IC50) values comparable to traditional biochemical assays. Notably, a model trained on cell line spheroids successfully classifies PDX spheroids, highlighting its adaptability.

Compared to convolutional neural network-based approaches, this method works with smaller training datasets and provides greater interpretability by identifying key morphological features. The droplet platform further reduces cell requirements, while single spheroid confinement enhances classification quality. Overall, this label-free experimental and analytical platform is confirmed as a scalable, efficient, and dynamic tool for drug screening.

## Introduction

One of the key challenges in oncology is the variability of responses to treatments and the frequent development of resistances to therapy.^1–3^ To tackle these complexities, research is continuously progressing in the understanding of molecular processes, the identification of new therapeutic targets and associated molecular markers, and the development of new drugs and therapeutic approaches. These findings are integrated into the general framework of “precision medicine”.^4^ Despite such progress, this strategy is not always sufficiently predictive, mainly due to tumor heterogeneity.^2,5^ A potential alternative or complementary approach would consist in testing drugs in vitro on patient-derived tumor models to identify the most suitable treatment options for each patient.^6^ This would facilitate the refinement of treatment selection on an individual patient basis (“personalized drug screening”). To be clinically relevant, this drug screening must be standardized, sensitive, and conducted in a short enough time to not delay patient treatment. In addition, it should be achievable with minimal cell quantities, to test a wide enough set of drug combinations and concentrations, using the limited number of cells provided by minimally invasive biopsies.^7^

The aim of refining treatment selection raises the question of the biological realism of the in vitro model. Conventional cell culture in monolayers at the bottom of culture wells (“2D culture”) was questioned because real tumors are 3D aggregates with potentially very different metabolic processes and permeability to drugs. Three-dimensional (3D) tumor spheroid models are thus preferred, for their ability to better mimic the heterogeneity, morphology, biological mechanisms, and the resistance of native tumors compared to traditional two-dimensional (2D) models.^8–11^

Finally, to allow screening of a reasonable variety of treatment conditions, current 2D or 3D in vitro cell culture methods most often require initial numbers of cells incompatible with most minimally invasive sampling methods. Microfluidic technologies have emerged as valuable tools to overcome this limitation, offering precise control over the cellular microenvironment while minimizing the consumption of cells and reagents in integrated systems.^12–15^ Various microfluidics systems have been developed to perform drug screening on 3D models^16^ using techniques like hanging drop culture,^17^ microwells,^18–23^ hydrogel encapsulation^24,25^ and droplet-based systems.^26–31^

However, using these models in personalized medicine presents challenges, especially when the gold-standard readout to evaluate cell viability after treatment is to conduct biochemical assays. Cellular assays such as live/dead fluorescent markers, for instance, offer precise results on cell viability, but they require extensive, costly and time-consuming image analysis, and are difficult to quantify in 3D.^32^ Additionally, many of these assays, such as MTT assay (a colorimetric assay for assessing cell metabolic activity) or live/dead, are endpoint assays and may lack precision in cytotoxicity measurement.^33,34^ Globally, biochemical assays are labor-intensive in both their implementation and analysis, which hinders their integration into routine treatment protocols.

The present work stems from the observation that exposed to cytotoxic agents, spheroids undergo specific morphological changes, as observed by us and others.^35,36^ These changes appear to contain a high amount of information, and this study aims to investigate whether machine learning methods can analyze and interpret these morphological changes, ultimately providing a label-free, quantitative, robust and reliable method for assessing the effectiveness of anticancer therapies. Some studies have already explored the use of deep learning for evaluating drug efficacy in tumor spheroids. For example, optical coherence tomography^37,38^ and refractive index tomography^39^ have been used to extract features from spheroid exposed to anticancer treatment, but these approaches required sophisticated and specific microscopy instruments and are difficult to scale up for high-throughput applications. Other approaches have used images of spheroids combined with deep learning algorithms for label-free analysis. Benning et al.^40^ and Tröndel et al.^41^ trained convolutional neural networks (CNNs) to classify spheroids into categories based on their response to drugs. Similarly, Chiang et al.^42^ developed a CNN model to classify spheroids and predict the drug concentrations to which they had been exposed to. Very detailed classification was achieved, e.g. as a function of the applied drug concentration, but to our knowledge, no prediction of drug efficiency was obtained. These studies, also, were conducted on cell lines only and focused on classification models that were trained directly on images.

This study adopts a streamlined and operational approach for the direct label-free prediction of drug efficiency, aiming for applications in clinical settings or drug development. This approach relies on the morphological analysis of spheroids using a machine learning model. Tumor spheroids were first generated and treated with drugs within droplets, using a previously described microfluidic platform.^43^ Brightfield images of the spheroids were recorded at several time points during treatment. To enhance image quality and enable automated imaging in controlled exposure conditions, a new cartridge was developed for use with a plate imager.

Unlike previous studies, however, this machine learning model was not trained directly on images, which can be sensitive to batch effect, e.g. between cell line and patient-derived specimen, and might not generalize well from small training sets. Instead, we opted for a two-step approach starting with a feature selection step before the classification task. To this end, a broad range of morphological features (e.g., size, texture,^44^ color) were first extracted from the images. Then, MIIC (Multivariate Information-based Inductive Causation) method^45,46^ was used to select the most informative features for classifying spheroids, as MIIC tends to outperform Deep-Learning methods in uncovering relevant information from small datasets.^47^ Finally, the most informative features were used to train a neural network to classify the spheroids into viability classes. As the ground truth for the machine learning model, a fluorescent end-point metabolic assay was performed to associate each spheroid with a discrete viability class, used for model training and accuracy assessment. The model was initially trained and tested on spheroids derived from a cell line, and then on spheroids derived from patient-derived xenografts (PDX). The study also evaluated the ability of a model trained on cell line spheroids to classify PDX spheroids and predict dose-response relationship. Using these classifications, “virtual” dose-response curves were generated, enabling the estimation of half maximal inhibitory concentration (IC50) values based solely on the morphological characteristics of spheroids. Then, to fully exploit the benefits of our platform, spheroid viability was estimated over the drug exposure time. Finally, the same label-free assay was applied on a dataset issued from another system to investigate the versatility of this approach.

These results demonstrate that the machine learning approach provides comparable insights to traditional biochemical assays, while leveraging a label-free and morphology-based methodology.

## Results

### The microfluidic platform facilitates spheroid production and drug treatment

Tumor spheroids were generated in droplets confined within tubing, using cells derived either from established cell line or PDX. The PDX samples were derived from Ewing Sarcoma, and the chosen cell line for this cancer was A673. Once in droplets, the cells in suspension aggregated spontaneously into a spheroid in less than 24 h.

First, growth and viability of the spheroids in droplets were assessed. In this purpose, PDX-derived spheroids were measured over a week and a metabolic assay was performed (see Method for details) to measure their metabolic activity over 8 days. Over this period, the spheroids grew continuously. This was confirmed with the increase in their metabolic activity, which doubled each day during the first 3 days. Then, it increased slower, and by day 8, the metabolic activity reached 800% of the level observed on day 1 (Figure S1).

After confirming that the spheroids were viable and growing during a period long enough for the drug assay (at least 3 days), a drug screening protocol in droplets was established (Figure 1A). Triplicates were performed systematically for both cell line and PDX.

**Figure 1.**
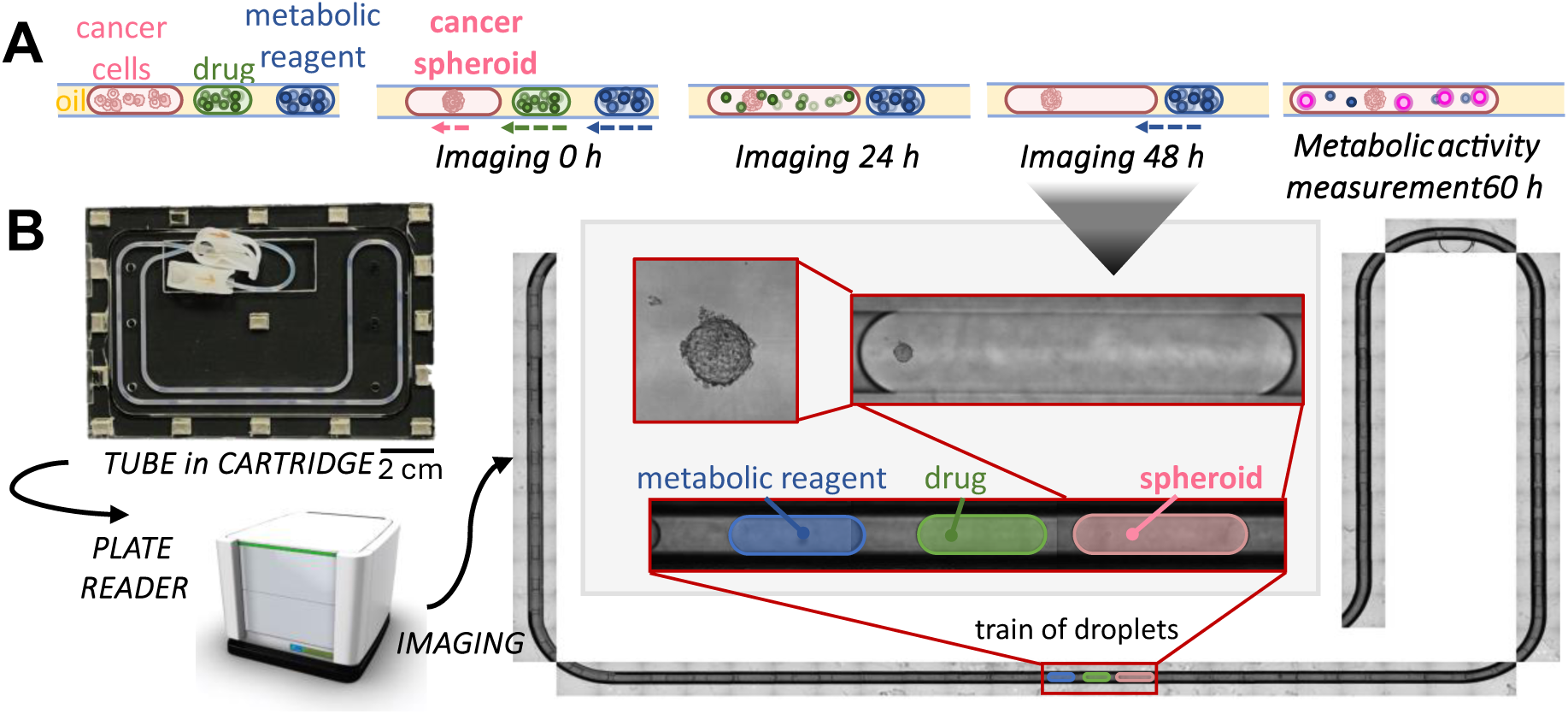
A) Workflow of drug testing in droplets. Trains of 3 droplets, with cells in suspension, drug solution and metabolic assay reagent are generated in tubes. After 24 h, the cells have formed a spheroid, and the drug droplet is merged with the spheroid droplet. Spheroids are exposed to the drug during 48 h. Then, as a ground truth for latter, the metabolic assay reagent droplet is merged with the spheroid treated with drug to measure its metabolic activity. The readout is performed after overnight incubation. Images are taken daily.

Imaging the spheroids inside droplets within tubes effectively presented a significant challenge. To address this, a custom-made device was designed to fit into a plate reader (Figure 1B). The device consisted of two micromachined cyclic olefin copolymer (COC) plates: one plate was transparent, allowing imaging through it, while the other plate featured a channel to securely hold the tube in place. The two plates were held together by magnets. To minimize optical aberrations, the same oil used to separate the droplets was introduced between the two plates. The plate reader was programmed to capture images along the length of the tube at different focal distances.

Images in brightfield were taken before drug exposure (0 h), after 24 h and after 48 h of drug exposure. A sample of images is presented for the spheroids derived from cell line Figure 2A and from PDX Figure 2B. In some droplets, several small spheroids were formed before drug addition, but they were fused the next day. It represented 7.5% of the cell line’s spheroids and 7.7% of the PDX spheroids. As observable on the images, the spheroids from both cell types undergo morphological changes depending on time and drug concentration. In the no-drug case, the spheroids increase in size. For increasing drug concentrations, a qualitative change in the morphology is observed between 0 h and 48 h, occurring between 2 and 5 µM for the cell line and between 1 and 2 µM for the PDX. The spheroids become darker, smaller and their texture change. This change of morphology is also visible after 24 h for spheroids submitted to the highest concentrations of drug. The main aim of this article is to check if this qualitative observation can be transformed, thanks to machine learning, into a quantitative, reproducible and predictive tool for drug response prediction.

**Figure 2.**
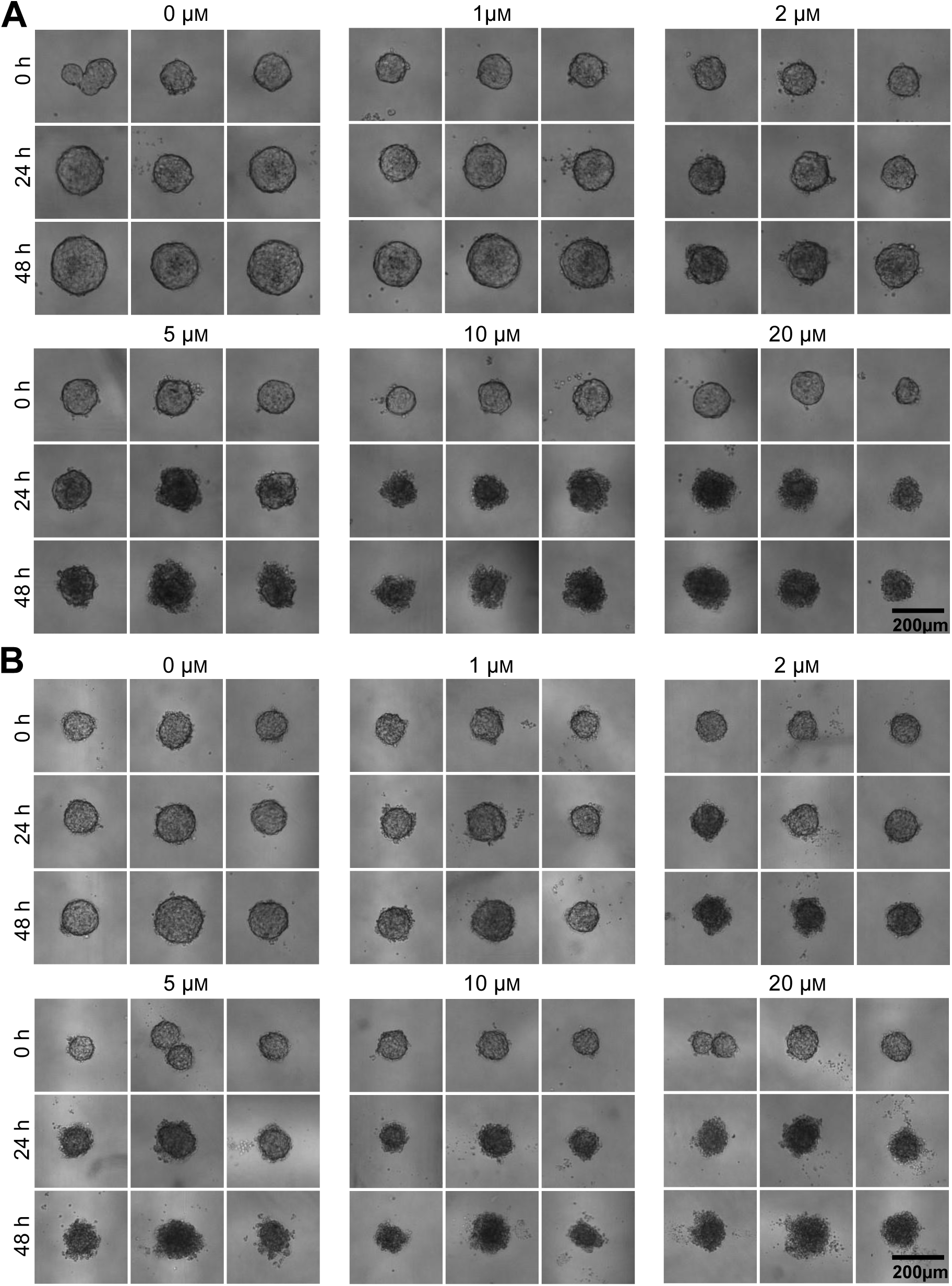
Images of spheroids in droplets derived from A) cell line and B) PDX cells, exposed to different drug concentrations (etoposide) after 0, 24 and 48 h of drug exposure. For each drug concentration, the same 3 spheroids are shown at different time points. Spheroids derived from cell line are cultivated from approximated 700 cells, and the ones from PDX from 1000 cells, in 1.8 µL droplets.

### Morphological features of spheroids evolve during drug treatment

After exposing the spheroids to a range of drug concentrations and imaging them at three time points, various morphological features were extracted, including area, circularity, mean grey value, as well as various texture descriptors (homogeneity, energy and correlation) derived from the computation of the Grey Level Co-Occurrence Matrix (GLCM).^48^ The full list of morphological descriptors is available Table S1. Since the microfluidic platform allows the tracking of each spheroid individually, it was possible to obtain the features of individual spheroids over time (Figure S2). For the (minority) of droplets containing multiple small spheroids, an averaged of the circularity, mean grey value and texture descriptors were taken while the area was summed according to the formula given in the Methods section.

The morphological features -- area, circularity, mean grey value, and correlation -- after 48 hours of drug exposure were then plotted depending on drug concentration and fitted to a sigmoid dose-response like curve. A selection of these plots is shown in Figure 3A for cell line and Figure 3B for PDX. Additional graphs for the features at 0 h and 24 h, as well as other texture descriptors and their variation, are presented in Figure S3 and S4 for the cell line and Figure S5 and S6 for the PDX. When possible, an IC50 value was derived from the sigmoidal fit, presented Figure S7. For the cell line, several indicators demonstrate concentration-dependent changes, including spheroid area, mean grey value and the correlation texture feature. In contrast, for the PDX model, only mean grey value could be used to fit a sigmoidal dose-response curve; and even for this feature the data points are more heterogeneous than for the cell line.

**Figure 3.**
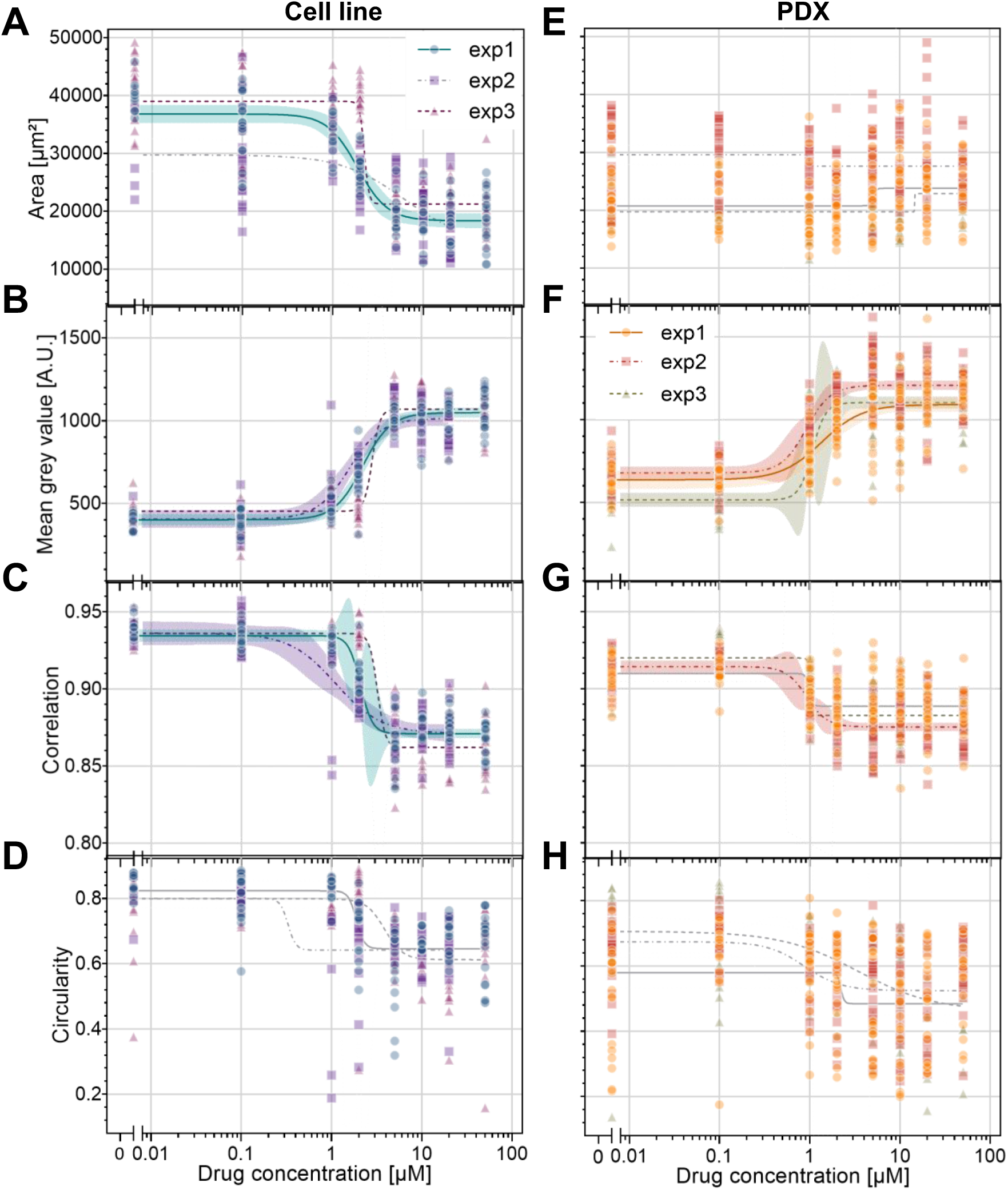
Morphological features of spheroids in droplet derived from A-D) cell line and E-F) PDX. Each combination of color and shape represents one independent experiment. Each point represents a single spheroid. The curves represent sigmoid fits with 95% confidence bands. The curves in grey represent curves with poor goodness-of-fit (R² < 0.5).

As an intermediate conclusion, morphological analysis alone seems to reveal significant trends upon drug concentration and time of exposure. However, it lacks the precision and reliability needed to accurately assess drug efficiency, especially for PDX, which are the most relevant to clinical situations. A supervised machine learning approach was thus developed to address this limitation. It first involved the establishment of a benchmark as a reference for training.

### Dose-response curve, using a metabolic assay, serves as a benchmark for assessing spheroid viability

As a benchmark for the machine learning approach, a metabolic bioassay was conducted after 48 h of drug exposure to evaluate the drug’s efficacy using a standard method. Each spheroid was then associated with an experimental metabolic activity score, defined as the fluorescence signal normalized to an averaged control (see Methods for details). Subsequently, dose-response curves were generated for each independent experiment, for both cell line (Figure 4A) and PDX (Figure 4B). Data were then fitted to a sigmoid curve, and the IC50 values were determined for each experiment. The average IC50 values obtained were IC_50,celle line_ = 3.2 ± 0.3 µM for the cell line and IC_50, PDX_ = 1.9 ± 0.2 µM for the PDX.

**Figure 4.**
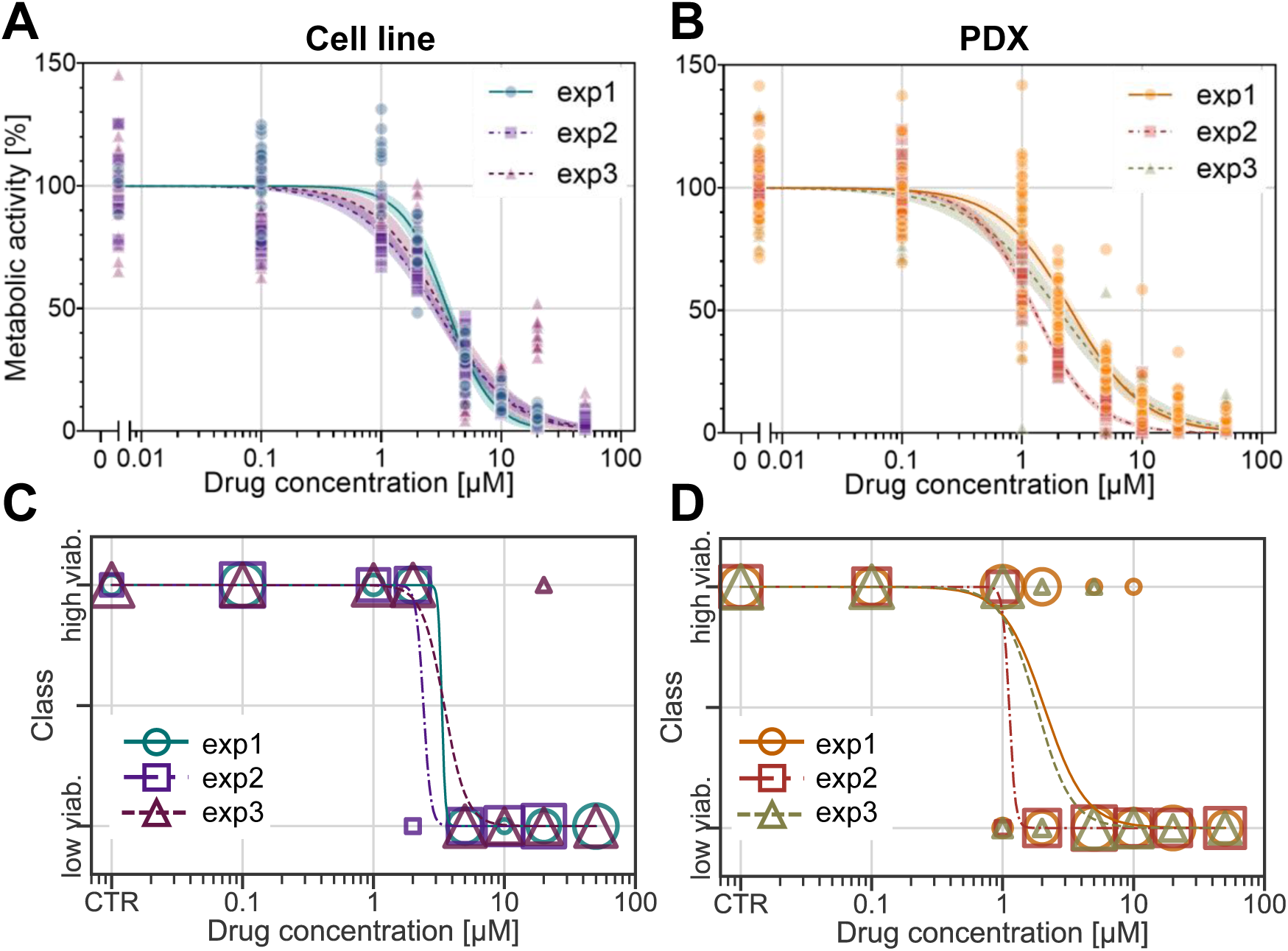
Dose-response curves with normalized metabolic activity depending on drug concentration (etoposide) of spheroids in droplets derived from A) cell line and B) PDX cells. C) And D) dose response curves with discretized metabolic activity into two viability classes depending on drug concentration for cell line and PDX respectively. Each combination of color and shape represents one independent experiment. For continuous metabolic activity, each point represents a single spheroid. For discretized metabolic activity, the size of each point on the plot is scaled proportionally to the number of overlapping data points. The curves represent sigmoid fits, with 95% confidence bands for the continuous metabolic activity.

This “bioassay-derived” metabolic activity score exhibits significant heterogeneity for a given drug concentration (e.g., for the cell line, the metabolic activity without drug ranges from 65% to 145%), reflecting the intrinsic uniqueness and variability of living systems. Moreover, this wide variability is also reflected in the morphological features of spheroids, where spheroids with similar metabolic activity display heterogeneous morphological properties (Figure S8). This variability poses a significant challenge to the algorithm’s ability to make meaningful predictions.

To address this issue, we used the experimentally observed fact that drug efficiency presents a relatively sharp threshold, as indicated by the sigmoid shape of the dose-response curves. On this basis, spheroids were classified as “low viability” for those with a metabolic activity below 50%, and “high viability” for those with a score above 50%. The dose-response curves generated after this classification are shown Figure 4C and 4D.

### The MIIC algorithm helps identify relevant morphological features

Our objective was to develop a machine learning-based approach that utilized spheroid morphological properties as input features and predict drug efficacy as the output.

To explore the relationship between morphological features and viability classes, the histograms and density distributions of spheroid features of the two cell types according to their assigned classes were plotted Figure 5A and 5C. Features for spheroids classified as “high viab.” and “low viab.” are shown in green and magenta, respectively.

**Figure 5.**
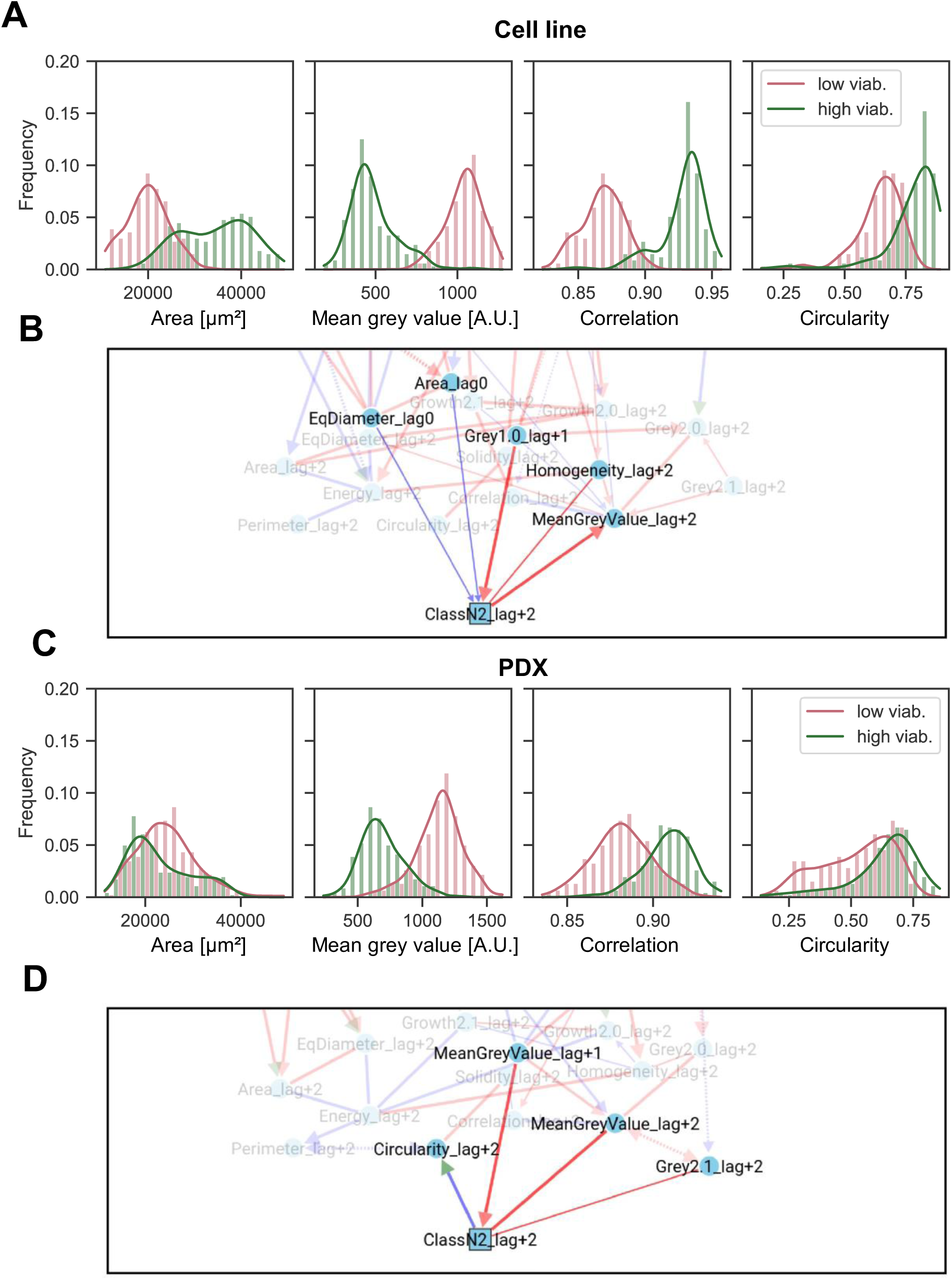
A) & C) Histograms of the distribution of 4 features after 48 h of drug exposure depending on their assigned class for respectively cell line and PDX. Curve represents the probability density for each class, bars show the normalized frequency of the data. N = 3.B) & D) Cropped MIIC network for respectively cell line and PDX.

For the cell line, the area, mean grey value and correlation graphs show minimal overlap between the two classes, suggesting that these features are robust features for classification. However, for the PDX, only the mean grey value graph presents classes with relatively low overlap, while the other plots show substantial overlap between the two classes. This is consistent with the observations in Figure 2, where this grey value seems to provide the most significant evolution with drug concentration. These curves, however, suggest that other features still contain some discrimination power, but provide no clue about the redundancy or independence of the contained information.

To identify the most relevant features for training the machine learning algorithm, the strength of association between each feature and the spheroid viability state was analyzed using the MIIC algorithm. It was applied to the dataset containing all extracted features for each spheroid, measured at different drug exposure times. This analysis computes the mutual information between each feature and the spheroid viability label, quantifying the feature’s importance in determining the spheroid class. Additionally, the algorithm retrieved associations between features. Some features are identified as having a direct possibly causal association with the spheroid class while other features are identified as having indirect impact on spheroid class. The features, their mutual information with the class and their relationship (direct or indirect) to the class are summarized in Table 1, and the networks highlighting the most informative features derived from the MIIC analysis are shown in Figure 5B and 5D for the two cell types. The full temporal networks are reported in Figure S9.

**Table 1.** The mutual information, in bits, shared between the class and the features. The column “direct link” indicates whether there is a link in the graphical network between class and the feature.

| Cell type | Feature | Direct link (Y/N) | Mutual information [bit] |
| --- | --- | --- | --- |
| Cell line | Homogeneity 48 h | Y | 1.360 |
| | $\Delta$ Grey Value 24 - 0 h | Y | 0.934 |
|  | Mean Grey Value 48 h | Y | 0.885 |
| | $\Delta$ Grey Value 48 - 0 h | N | 0.862 |
|  | Correlation 48 h | N | 0.816 |
|  | Growth 48- 0 h | N | 0.786 |
|  | Diameter 48 h | N | 0.641 |
|  | Area 48 h | N | 0.641 |
|  | Energy 48 h | N | 0.636 |
|  | Solidity 48 h | N | 0.540 |
|  | Circularity 48 h | N | 0.498 |
|  | Perimeter 48 h | N | 0.281 |
|  | Diameter 0 h | Y | 0.028 |
|  | Area 0 h | Y | 0.028 |
| PDX | Mean Grey Value 24 h | Y | 0.646 |
|  | Mean Grey Value 48 h | Y | 0.617 |
| | $\Delta$ Grey Value 48- 0 h | N | 0.621 |
| | $\Delta$ Grey Value 24 - 0h | N | 0.562 |
|  | Correlation 48 h | N | 0.376 |
|  | Correlation 24 h | N | 0.371 |
|  | Solidity 48 h | N | 0.165 |
|  | Circularity 48 h | Y | 0.155 |
|  | Circularity 24 h | N | 0.101 |
|  | Perimeter 48 h | N | 0.079 |
|  | Solidity 24 h | N | 0.069 |
| | $\Delta$ Grey Value 48 - 24 h | Y | 0.065 |
|  | Perimeter 24 h | N | 0.061 |
|  | Area 48 h | N | 0.043 |
|  | Diameter 48 h | N | 0.043 |

The MIIC network also enabled the identification of more subtle relationships between features. For the cell line, the initial spheroid size (prior to drug exposure) showed a slight influence on the final classification in terms of mutual information. This suggests that smaller spheroids derived from this cell line may exhibit slightly greater sensitivity to the drug compared to larger ones. Additionally, the presence of multiple small spheroids before drug exposure was found to have no measurable impact on spheroid classification, justifying our initial choice of integrating them into the dataset.

The features identified by MIIC as the most critical for spheroid classification are consistent with the qualitative intuitions derived from classes overlap. These features display an important variation across classes, but their specific or redundant information about classes is not straightforward to assess. In this context, MIIC analysis offers a quantitative evaluation of each parameter’s significance, providing deeper insights into their specific contributions to spheroid classification.

### A supervised machine learning model classifies spheroids in viability classes based on their morphological features for IC_50_ prediction

A supervised machine learning model based on a neural network was then trained to classify the spheroids into two classes, “high viability” and “low viability”. Input features were selected based on their relevance, as identified by the MIIC algorithm. The features selected to train the model were the 15 ones with the highest weight. For the cell line, as data at 24 h was missing for one of the experiments, only features at 0 and 48 h were selected.

The algorithm was trained on 2 out of the 3 experiments and tested it on the 3^rd^ experiment. The process was repeated for each experiment as a cross-validation process. The detailed number of points in each class and each experiment is detailed Figure S10A and S10B. To evaluate the performances of the model, the ROC-AUC scores are presented in Figure 6A and 6D as well as four metrics; accuracy, precision, recall and f1-score, presented in Figure 6B and 6E. Essentially, the closer these metrics are to 1, the better the classification performance. For the cell line dataset, the model achieved near-perfect classification, with a f1-score of 96%. For the PDX, the model performance was slightly lower (f1-score of 93%). The curves representing the accuracy VS epoch are visible Figure S11, indicating that the model did not overfit.

**Figure 6.**
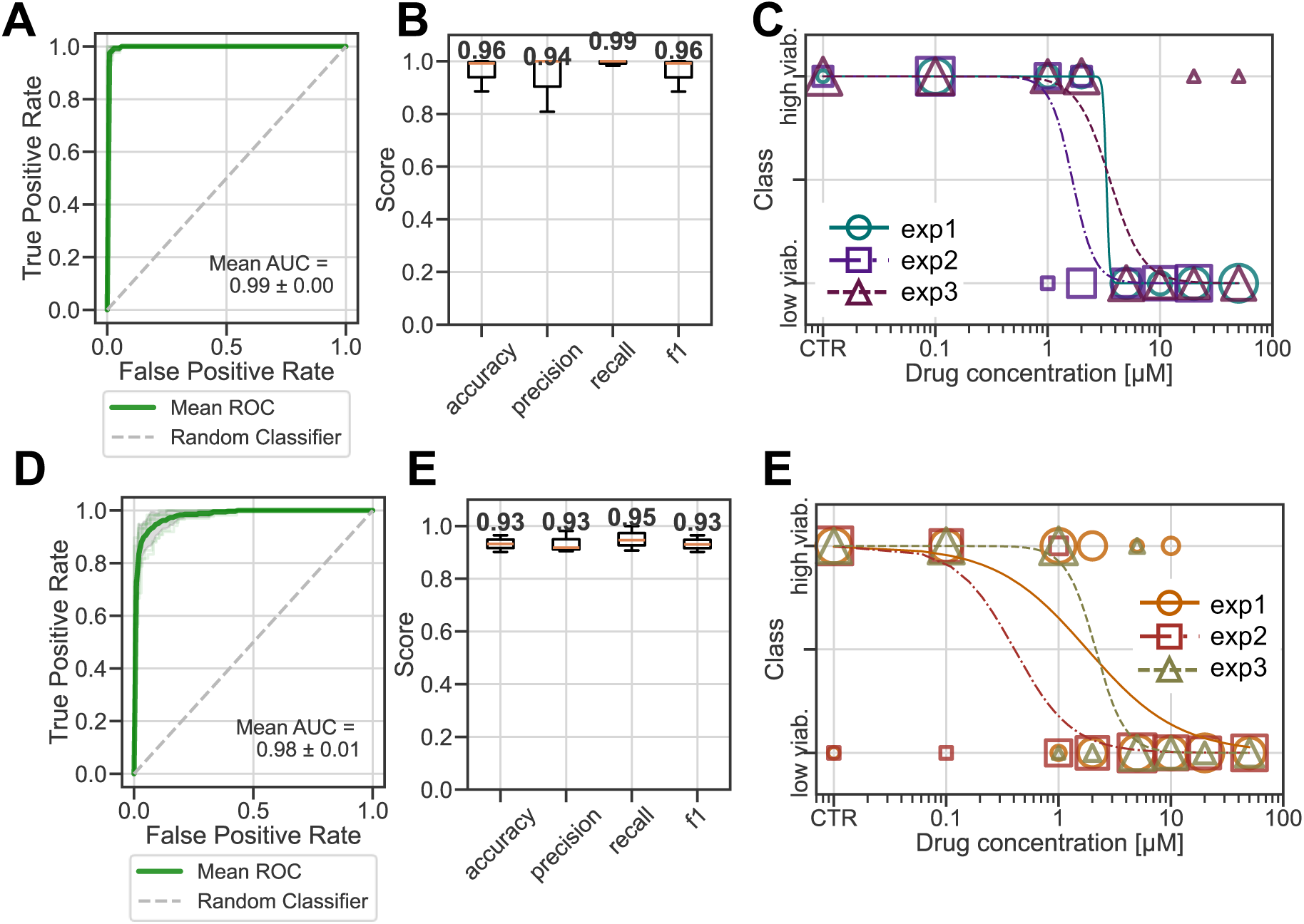
Results of machine learning for A-C) cell line and D-E) PDX. A&D) ROC curves. B&D) averaged metrics. C&E) Dose-response like curves after classification. Each combination of color and shape represents one independent experiment. The size of each point on the plot is scaled proportionally to the number of overlapping data points.

This classification was then used to estimate drug efficacy. To this end, the predicted class for each spheroid (0 for “low viab.” and 1 for “high viab.”) was plotted as a function of the drug concentration to which the spheroid was exposed, similar to a dose-response curve (Figure 6C and 6F). The comparison between the IC50 values obtained using the machine learning approach and the metabolic assay approach is shown in Figure 7 and Table 2. Indeed, one can see that the machine learning method provides an IC50 value close to that of the metabolic assay used as an initial reference.

**Figure 7.**
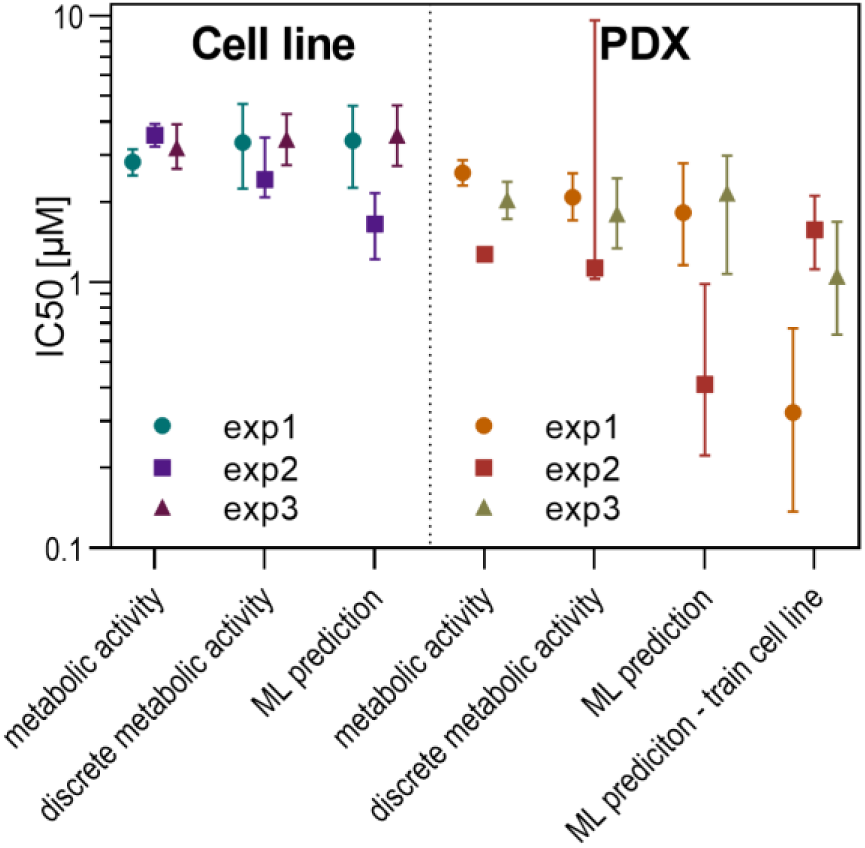
IC50 values computed from metabolic activity and inferred from machine learning classification for cell line and PDX. Each point represents an independent experiment. The error bars represent the 95% CI.

**Table 2.** IC50 values for cell line and PDX computed with different methods.

| Cell type | Assay | IC50 [μM] | | |
| --- | --- | --- | --- | --- |
|  |  | exp1 | exp2 | exp3 |
| Cell line | Metabolic activity | 2.8 | 3.6 | 3.2 |
|  | Discrete metabolic activity | 3.3 | 2.4 | 3.4 |
|  | Machine learning prediction | 3.3 | 1.7 | 3.6 |
| PDX | Metabolic activity | 2.6 | 1.3 | 2.0 |
|  | Discrete metabolic activity | 2.1 | 1.1 | 1.8 |
|  | Machine learning prediction | 1.8 | 0.4 | 2.1 |
|  | ML prediction, training cell line | 0.3 | 1.6 | 1.1 |

To assess the robustness of our classification into two viability classes, the same procedure was repeated using a “three classes” classification. Spheroids were classified as “low viab.” if their score was below 25%, “high viab.” if it was above 75%, and “intermediate” for scores in between. The distribution of these classes is shown in Figure S10 C&D. The results regarding viability and IC50 did not differ significantly from those obtained with the two-class model, despite lower performance metrics, with a f1-score of 64% for the cell line and 75% for the PDX model (Figure S12). This supported our initial decision to use a binary classification model.

One of the key advantages of the droplet platform is its ability to track spheroids over time at a single-scale level, enabling dynamic assessments of spheroid viability during drug exposure. To explore this potential, the neural network model was trained, as before, to classify spheroids based on their viability class. However, this time, only the features measured after 48 hours were used to train the model. Then, the features at 24h were used to make predictions on spheroid viability class. From this classification, dose-response curves and IC50 values were inferred (Figure S13 and Table S3).

The comparison of IC50 values inferred from morphological features at 24 hours with those obtained from the metabolic assay at 48 hours revealed, for the cell line, no significant differences. However, for the PDX, a slight shift of IC50 toward higher concentrations was detected, as expected for spheroids exposed to the drug for a shorter time. These results highlight the platform’s potential for dynamic drug sensitivity assessments over time. However, since the metabolic activity scores were only known after 48 hours of drug exposure, these predictions could not be validated against a definitive ground truth.

### A model trained on cell line data successfully predicts PDX response

Finally, to check the robustness and generality of the approach, targeting clinical use, the neural network model was trained using data derived from cell line spheroids and subsequently tested on PDX spheroids to evaluate its predictive accuracy in a different biological context. The set of features that were identified as important for the PDX were used.

After applying the trained model to the PDX spheroids, the overall accuracy of the predictions was equivalent to the one obtained previously, when training and testing on the PDX dataset. The model’s performance is highly satisfactory, with a classification accuracy of 87%, demonstrating a generalization capacity across different sample types (Figure 8).

**Figure 8.**
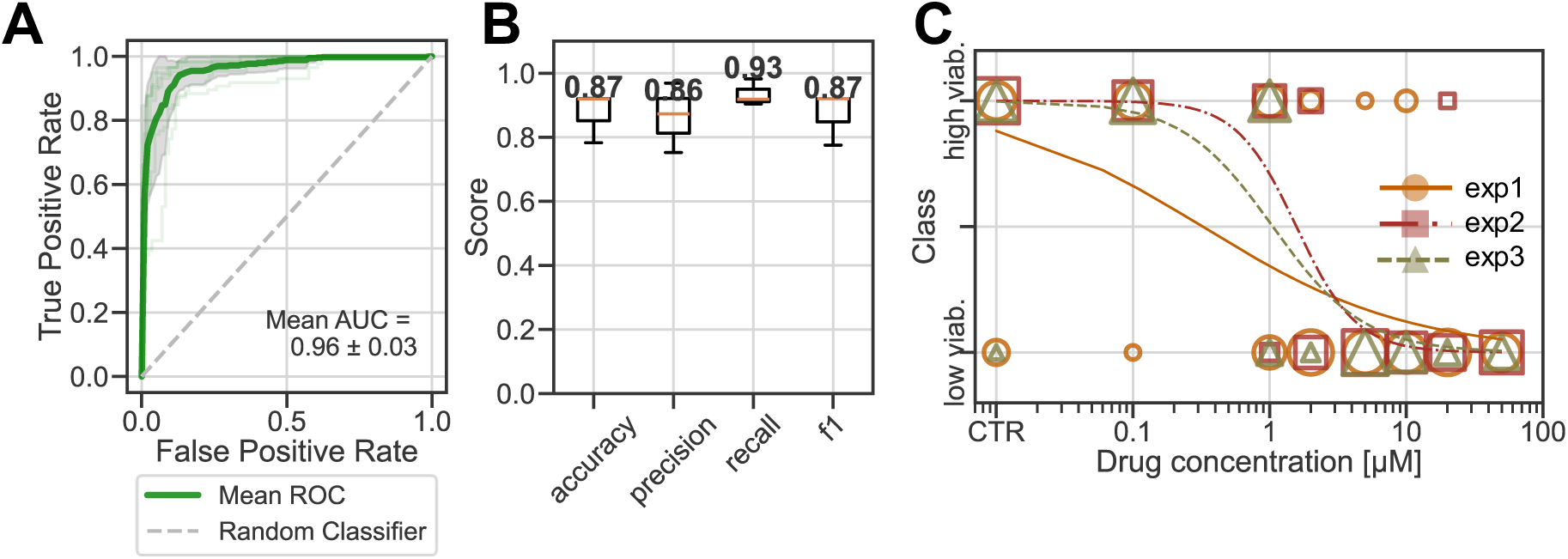
Results of machine learning algorithm trained on cell line dataset et tested on PDX dataset. A) ROC curve, B) metrics and C) dose-response curve after classification. Each combination of color and shape represents one independent experiment. The size of each point on the plot is scaled proportionally to the number of overlapping data points.

Moreover, the model demonstrated an ability to predict the drug response profiles of PDX spheroids, identifying patterns of resistance and sensitivity that aligned closely with those observed through biochemical assays. This suggests that despite the inherent biological differences between cell lines and PDX samples, the features learned by the neural network from the cell line data could retain relevance in the context of patient-derived samples.

## Discussion

This study advances the framework of a droplet-based microfluidic platform for anticancer drug screening in 3D by incorporating a label-free assay to assess drug efficiency. The long-term objective is to provide a platform allowing fast, high-throughput and label-free screening of anticancer drugs on tumour spheroids, using a minimal number of cells.

The first challenge encountered was the integration of in situ and dynamic brightfield imaging. To address this, a custom tube holder device was designed to be compatible with a commercial automated plate reader, allowing spheroid imaging directly within droplets. This approach has the advantage of using commonly used laboratory equipment. However, it requires an operator to manually launch image acquisition for each tube. Additionally, the plate reader, designed to image evenly spaced wells, captured many unnecessary images without spheroids, requiring pre-processing of images before analysis. A (time-consuming) manual step was set to select the spheroids among the images, followed by an automated spheroid segmentation. While a future integration of AI for automated spheroid identification could significantly reduce analysis time, we opted for manual selection at this stage to ensure precise identification, particularly for spheroids at the edges of droplets or slightly out of focus. Not all images were usable due to these limitations, but manual sorting ensured reliable tracking of the spheroids. Further development in imaging and tube-holder design specifically tailored for this droplets-in-tubes platform will be to fully automate both imaging and analysis, reducing hands-on time, increasing throughput, to enhance the platform’s applicability for pharmaceutical research and clinical applications.

The images obtained from both the cell line and the PDX spheroids treated with drug suggested at first glance a qualitative correlation between their appearance and the amount of drug they have been exposed to. The quantitative analysis revealed that area, mean grey value and texture correlation varied with drug concentration in cell line. However, for the PDX model, the data were more heterogeneous, with only mean grey value showing a significant variation in response to drug concentration. These observations led us to explore a more systematic and robust approach to assess drug efficacy based on morphological features.

In this purpose, we developed a supervised machine learning model to provide quantitative label-free assessments of drugs efficiency. We aimed at minimizing the size of the dataset and the computing power needed for training and for operation. For that, unlike prior approaches directly training the model on images, we chose a two-steps approach, in which various morphological descriptors are first obtained by conventional image analysis and then used in a supervised machine learning approach. Supervised training was based on a metabolic assay that associates a metabolic activity score to each spheroid. To avoid the dispersion associated with the use of a continuous metabolic score variable, and its detrimental effect on the size of the dataset required for training, each spheroid was classified as either ‘low viability’ or ‘high viability’ based on its metabolic activity score. Each spheroid is then associated with its morphological properties at different drug exposure times and with its viability class.

To better understand how various morphological features contribute to the spheroid’s classification, we used a causal network learning algorithm, MIIC, to identify the relationships between the features and their relative importance for classification. After identifying the 15 more relevant features ranked by the MIIC, we trained a neural network model using these features to predict spheroid viability class. The model was first trained and tested on a cell line dataset and then on a PDX dataset, both from Ewing Sarcoma.

Our model successfully classified spheroids into two classes with an accuracy of 96% and 93% with dataset of 316 and 428 data points for cell line and PDX, respectively. Other machine learning models were also tested, showing similar performance (Figure S14). This accuracy is higher than the one reported in Benning et al.^40^ (90%) and close to the one reported by Tröndle et al.^41^ (98.2%), However, the performance was achieved here with a smaller dataset, in contrast to direct deep learning from images, which are typically trained on larger image datasets (e.g. for instance n = 4974 for Tröndle et al., n = 1200 for Benning et al.). The ability to use a relatively small dataset makes this approach more suitable for clinical applications for precision medicine, where the number of available cells from biopsy, and thus the number of possible tests, is often limited.

Importantly, the label-free assay developed in this work has the potential to be adapted for broader applications beyond this platform and integrated into other systems. To evaluate the approach’s independence from the experimental method, the algorithm was trained and tested on spheroids formed in agarose microwells (see Figure S15 and S16 and supplementary Methods). The classification f1-score reached 89% for cell line and 81% for PDX (Figure S17). Although this score is lower than that achieved in droplets, it remains highly acceptable. This demonstrates that the strategy, based on features extraction and neural network classification, can be applied to other experimental platforms, provided they produce high-quality images of individual spheroids. However, the final accuracy is platform dependent. Systems such as the droplet platform, with high spheroid-to-spheroid reproducibility in terms of growth, drug exposure, and screening conditions, are expected to yield better results, as shown in this proof of concept.

Remarkably, the reliable classification achieved when training on the cell line dataset and testing on the PDX dataset opens up the possibility of ultimately predicting drug efficacy in primary patient-derived cells using a label-free approach. To further validate this objective, the next steps involve expanding the approach by training models of a broader range of PDX spheroids across different cancer types and assessing that they share sufficiently similar morphological properties.

Finally, the predicted class of each spheroid was plotted against drug concentration, and a sigmoid curve was fitted to compute a “morphological” IC50 value. The comparison of IC50 values derived from the metabolic assay and the machine learning model yielded consistent results. Misclassified spheroids were typically those exposed to drug concentrations near the IC50, where an intermediate state might exist, with dying spheroids not yet exhibiting the same features as fully dead ones. We explored this hypothesis by adding an intermediate state between ‘alive’ and ‘dead’, but the results showed poor classification accuracy. This may be explained by the steep decrease in metabolic activity around the IC50 in our dataset. In cancers with a less sharp drug response, a three-class classification might better capture the spheroid states.

It should be noted that in vitro assays serve only as indicators of drug efficiency, so that their transposition to clinical settings requires an adaptation for each specific situation.

While the combination of this label-free assay and our droplet platform demonstrates strong synergy, we believe that this method holds the potential for broader application to datasets generated using other systems. For example, we successfully applied the same label-free classification approach to images of spheroids grown, treated, and imaged in an agarose microwell system, achieving reliable accuracy. This highlights the versatility of our method for use in various experimental settings.

## Conclusion

This study demonstrates the potential of machine learning in developing a label-free assay for evaluating drug efficacy in cancer spheroids. First, spheroid morphology was shown to correlate with viability after chemotherapy treatment. However, direct inference of drug efficiency from these observations lacks robustness and consistency across datasets. To address this, we developed a machine learning approach, able to better capture information embedded in spheroid appearance. We extracted morphological features such as size and color from brightfield images, selected the most relevant ones using MIIC model and used these key features to train a neural network to classify the spheroids based on their viability before generating drug-response curves and IC50 estimations.

This approach offers a simple and robust method for assessing drug treatment efficacy. It outperforms or matches approaches based on deep learning models directly trained on spheroid images, while using less than 500 spheroids. Additionally, it allows for a deeper understanding of the morphological features driving spheroid classification. This insight paves the way for its broad applicability across different cancer types or drug screening systems, using the same machine learning model for diverse datasets.

The integration of this label-free approach with a droplet-based microfluidic platform offers a powerful tool for drug screening. The droplet platform ensures reproducible spheroid generation and testing conditions while minimizing cells - key advantage for rare cancers, minimally invasive biopsies, large-scale drug screening, and routine clinical workflow. Furthermore, its compatibility with plate readers ensures consistent imaging conditions.

In summary, this approach combines simplicity, efficiency, and broad applicability, offering significant potential for drug screening in cancer research and clinical application.

## Materials & methods

### Cell line culture

The Ewing sarcoma A673 cell line (ATCC CRL-1598) was cultured at 37°C and 5% CO_2_ in Dulbecco’s modified Eagle’s medium (DMEM, Gibco^TM^, 61965-026), supplemented with 10% fetal bovine serum (FBS, Dutscher, S1900-500C) and 1% penicillin/streptomycin (Gibco^TM^, 15140-122). Mycoplasma testing was conducted every 1 to 2 months.

### Patient-derived Xenograft cells

Patient-derived xenografts (PDX) of Ewing Sarcoma (IC-pPDX-87) were provided by INSERM U1330 (authorization APAFIS #43745-2023060615213570 v2, 24/06/2023 given by National Authority). The tumors were dissected and dissociated to obtain cell suspension of cancer cells. The dissociation was performed as previously described in Buchou et al.,^49^ in RPMI (Sigma, R8758) using 150 µg/mL Liberase (Roche, 5401020001) and 150 µg/mL DNase (Sigma, DN25-100MG), for 30 min at 37 °C with gentle mixing. Cellular viability was quantified using a Vi-cell XR Viability Analyzer (Beckman Coulter, Brea, CA, USA). Once collected, the cells were diluted at the desired concentration in DMEM-F12 (Gibco^TM^, 31331-028) supplemented with 2% B27 (Gibco^TM^, 17504044) and 1% Penicillin/Streptomycin; and used in the few hours following the dissection.

### Droplet microfluidics platform

The droplet microfluidic platform used for generating and culturing tumor spheroids, as well as for conducting drug screening, is detailed in Parent et al..^43^ Briefly, droplets were formed within PTFE tubing (Adteck Polymer Engineering^TM,^ BIOBLOCK/14) of internal diameter 0.81 mm and 50 cm long. The droplets were automatically aspirated using a syringe pump (Tecan Systems, Cavro® XMP 6000) that pipet the different solutions from a well plate into the tubes. The system was capable of simultaneously producing 8 tubes, each containing 20 trains of 3 droplets, with each tube corresponding to a different drug concentration.

Each droplet train consisted of one cell suspension droplet (1.8 µL), one drug solution droplet (1.2 µL), and one metabolic activity assay droplet (1.2 µL), separated by oil phase containing surfactant (Fluigent, dSurf) with volumes of 0.45 µL, 0.9 µL respectively, and 1.0 µL between two trains of droplet. The working oil was FC-40 oil (Merk, F9755), present at the tube extremities. The biocompatible oil (dSurf) was separated from the FC-40 oil with a droplet of PBS (Sigma-Aldrich, D8537) of 6 µL. After droplets formation, the tubes were clamped at both ends and incubated in a black box to protect them from light.

The droplets can be merged on demand to implement protocols with several steps of reagents addition. The merging principle is detailed in Parent et al.^43^ and Ferraro et al.^50^ Briefly, at a sufficiently high flow rate, smaller droplets move faster than larger ones, enabling the adjacent droplets to get closer together and then to merge them. In practice, the tubes were plugged back onto the syringe pumps. Then, the droplets were moved at 6 µL.s^-1^ for 45 µL, then brought back to the initial position at 0.3µL.s^-1^. This process was repeated to achieve droplet-to droplet contact, and merging was induced by an electric field. As previously discussed,^43^ some droplet pairs occasionally failed to merge. These unmerged droplets were easily identified in the imaging process and discarded from the analysis.

### Spheroid production, drug treatment and drug efficacy readout in droplet

The droplets in tubes were generated as described before. For experiments with cell line A673, cells were diluted at 400 000 cells.mL^-1^ and for experiments with PDX, at 550 000 cells.mL^-1^ to reach approximately 700 cells and 1000 cells per droplet, respectively. The drug, etoposide (GreenPharma, Prestw-396), was diluted in the same culture media as the cells (but not supplemented for the PDX experiment) at 7 different concentrations between 0.1 and 50 µM. The metabolic assay, Alamar Blue (Invitrogen^TM^, A50101), was diluted in cell culture media to reach a final concentration of 20% (%v/v) after droplet merging.

After 24 h of culture, spheroids were treated with the drug by merging the droplet containing the spheroids with an adjacent droplet containing the drug. After 48 h of treatment, the droplets containing the metabolic assay were merged with those containing the drug-treated spheroids.

For both cell line and PDX, 3 independent experiments were performed at different times. For the cell line experiment, one experiment consisted in testing 8 conditions (no drug and 7 drug concentrations) with a maximum of 20 replicates per condition, depending on the droplet merging. For the PDX experiment, one experiment consisted in testing 8 conditions two times, so a maximum of 40 replicates per condition.

### Metabolic activity measurement for spheroid viability assessment

After one night of exposure to the metabolic assay, a readout was performed. AlamarBlue is based on resazurin, which is reduced into a fluorescent species in contact with viable cells, leading to an increase of the droplet fluorescence depending on the metabolic activity of the spheroid inside this droplet. The fluorescence of all the tubes were measured with a scanner (Typhoon FLA 9000, filter Cy3, photomultiplier value of 250 V, pixel size of 25 µm).

For each independent experiment, the average fluorescent intensity of each droplet was calculated from the fluorescent image using a python script. It was then normalized by subtracting the basal response and dividing by the mean fluorescence of the control droplets (without drug treatment) to obtain a normalized metabolic activity value, as follow in Equation 1:

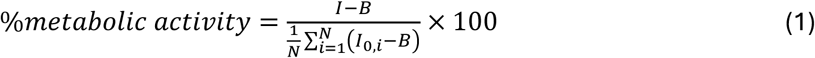

Were *B* corresponded to the basal signal of Alamar alone (without cells), *I_0,i_* to the fluorescence intensity of a control droplet (no drug case) and *I* to the fluorescence intensity of the droplet.

The dose-response curve for each experiment was plotted with GraphPadPrism (version 9.3.1) and a sigmoid fit was performed using an [inhibitor] vs. normalized response with a variable slope, corresponding to the Equation 2:

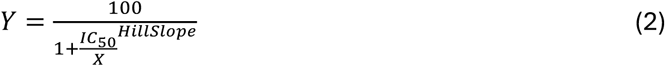

Where IC_50_ and Hill Slope are features estimated from the fit, and *X* are the values of metabolic activity.

### Image acquisition

Images of the spheroids were taken with a plate reader (Perkin Elmer, EnSight®), equipped with an objective 4X. Each tube was installed into a home-made device consisting of two cyclic olefin copolymer (COC) plates. The two plates were maintained together with 17 pairs of magnets (Supermagnete, Q-06-04-02-HN and Q-07-06-1.2-N) distributed along the plates. The tube was inserted into the designed plate (Figure S18) and HFE oil (Fluorochem, F051243) was inserted between the two plates. The plate reader was programmed to take images along the tube in different focal distances. Images were taken every 25 µm, in brightfield, and the parameters of illumination were fixed to 6% of power and 6 ms of exposure time.

### Image analysis and spheroid segmentation

As the imaging system took pictures of the entire tube, many of the images did not contain spheroids. To address this, a custom program was developed to allow users to quickly identify the locations of the spheroids. The program then cropped the image stack around the spheroid and selected the best image in the focal plane by choosing the image with the highest sharpness, given by the variance of the Laplacian. Segmentation was performed using the Canny Edge algorithm. A binary mask for each spheroid was extracted from which various features were computed: area, circularity, as well as colorimetric and texture features.

The area was calculated as the number of pixels contained within the mask and converted to µm² using the image resolution (1 pixel = 3.26 µm). Circularity was determined using the Equation 3:

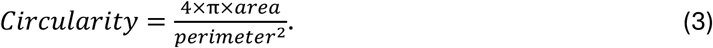

Mean grey value was measured within the spheroid, excluding the edges of the spheroids, and the background value was subtracted. The background value was computed by averaging pixel values outside the spheroid mask.

Texture features were extracted using Gray-Level Co-occurrence Matrix (GLCM) analysis,^44,48^ focusing on three key parameters: homogeneity, energy, and correlation. GLCM is a statistical tool that quantifies the spatial relationships between pixel intensities, providing insights into the underlying structural patterns of the texture. Homogeneity indicates the closeness of GLCM elements to the diagonal, reflecting texture uniformity—higher values represent smoother textures. Energy (also known as Angular Second Moment) evaluates the sum of squared GLCM elements, representing texture uniformity and increasing in images with repetitive or regular patterns. Correlation measures the linear dependence between pixel values at specified offsets, capturing the complexity of texture through pixel similarity with neighboring pixels.

The evolution of some parameters over time was also studied, by computing the difference of values before and after drug exposure. Some droplets before drug addition contained several small spheroids that fused latter. In these cases, the spheroid properties were either averaged (for textural, colorimetric parameters, and circularity) or combined for area and diameter using Equation 4:

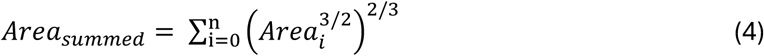

### MIIC

The MIIC algorithm is a network reconstruction method to analyze large scale biological or biomedical data,^46^ which has recently been extended to analyze morphodynamic features extracted from time-lapse images of cellular systems (Simon 2025). In the present study, this temporal algorithm (tMIIC) was used as a feature selection approach to identify the most informative morphological features to predict the viability of the spheroids after 48 h in the presence of drug. MIIC aims to learn a large class of causal and non-causal graphical models, including the effects of unobserved latent variables, corresponding to the presence of an unobserved common causes and represented by a bidirected dashed edge. The non stationary temporal version of the algorithm tMIIC was used to reconstruct the temporal network starting from the spheroids temporal data. The algorithm can handle missing data with no need of imputation. The number of temporal layers was set to 3 as each temporal layer corresponds to a different day of image acquisition and feature extraction. The algorithm outputs a network, Figure S9, where each node represents a variable and each edge encodes for a dependency possibly lagged between the 3 time points (0 h, 24 h and 48 h). An edge with an arrow stands for causal relations, while an edge without arrow stands for association (*i.e.* its causal or non-causal nature cannot be determined from the available data). Green arrows stand for genuine cause-effect relations while an arrow that is not green stands for a putative causal effect, which can either be a genuine causal relation or the effect of a latent variable (*i.e.* a bidirected edge), although this cannot be determined from the available data (Ribeiro-Dantas 2024). The colors of the edges are based on the Spearman’s correlation coefficient computed for the two linked variables. The color red stands for positive correlation, the color blue for negative correlation. The variables listed in Table1 are those sharing the highest mutual information measured in bits. The letter “Y” in “Direct link” column corresponds to direct effect, as the Class and the features are linked in the graph. The letter “N” corresponds to indirect effect, as the Class and the features are not directly linked but they are significantly associated in terms of shared mutual information. A Python script was used to prepare the data in the format needed by the MIIC algorithm. The MIIC method used is implemented in R.

### Machine learning model

A neural network model was created to assess a supervised classification task. The input were the morphological features of the spheroids at different time points. The target was the class of the spheroid, encoded from its metabolic activity score.

The neural network model was implemented in Python (v3.12.7) using Keras library. It consisted of 3 dense layers, starting with an input layer that accepted vectors of dimensionality matching the size of the input data. The first hidden layer comprised 128 units and employed the rectified linear unit (ReLU) activation function to introduce non-linearity, allowing the model to capture complex patterns in the data. This was followed by a second hidden layer with 10 units, also using the ReLU activation function, designed to further reduce the dimensionality of the input while preserving important features for classification. The output layer contained a number of neurons corresponding to the total number of target classes, and a softmax activation function was applied to produce a probability distribution over these classes. The model was compiled using the Adam optimizer. The loss function was set to categorical cross-entropy, which is well-suited for classification tasks involving mutually exclusive classes. The model’s performance was tracked using accuracy as the primary metric, enabling the assessment of its classification performance during training and validation.

As each dataset (cell line and PDX) was composed of 3 independent experiments, a cross-validation approach was used where the model was trained on 2 independent experiments and tested of the third experiment, and so on for each of the 3 experiments.

Spheroids with missing data for one or more features were excluded from the dataset. For the A673 cell line, the dataset was composed of 316 spheroids, and for the PDX of 428 spheroids. The detailed dataset descriptions are provided in the supplementary material (Figure S10).

To assess the performance of the model, different metrics were measured for each training/testing set. First, the Receiver Operating Characteristics (ROC) curve was computed and its area under the curve (ROC-AUC) measured. Then, the accuracy, the precision, the recall and the f1-score were computed to get insight on the model performance. Finally, during training, accuracy and loss were computed at each epoch for both the training and test datasets. These metrics were monitored to assess model performance and detect overfitting.

### Inference of the IC50 from the machine learning class prediction

For each independent experiment, after the classification of the spheroids with the machine learning model, the predicted class of each spheroid depending on the drug concentration it has been exposed to was plotted. A sigmoid curve with a variable slope, constrained to plateau between 0 and 1, was fitted on these points and an IC50 value was inferred for each experiment.

A Bayesian inference approach was used to estimate the IC50 and slope parameters of the dose-response curve, using the Markov Chain Monte Carlo (MCMC) sampler from the Python package emcee.^51^ A sigmoid model was fitted to the points. IC50 and slope were assigned uniform priors within the ranges 0–20 and −50 to −0.1, respectively. The likelihood function was defined based on a logistic model for binary classification. The MCMC sampler was initialized with 32 walkers and run for 500 iterations following a burn-in phase, generating posterior distributions for the parameters. Median parameter values and 95% credible intervals were calculated from the posterior samples.

## Supporting Information

Supporting Information is available from the Wiley Online Library or from the author.

## Acknowledgements

This work was supported in part by ANR PRCE DROMOS (ANR-20-CE19-0012) and by the European Research Council funding (ERC-2019-COG NanoBioMade-865629). CP acknowledge a PhD fellowship PSL-Qlife (ANR-17-CONV-0005). This work has benefited from the technical contribution of the joint service unit CNRS UAR 3750. The authors would like to thank the engineers of this unit for their advice during the development of the experiments. CP and HH contributed equally to this work.

## Conflicts of Interest

The authors declare no conflict of interest.

## Author Contributions

CP and HH performed the experiments, analyzed the data and implemented the algorithms. CP developed experimental protocols and methodology. TT, FS and HI performed the MIIC analysis. AJ created the device for imaging. SZ provided PDX cells. OD, CW and JLV conceptualized the project. CP, HH and JLV wrote the initial draft. All the authors reviewed and approved the manuscript.

## Data Availability Statement

The codes and raw data used in this study are openly available in GitHub repository at https://github.com/cparent3/spheroid_morpho_ML. The data that support the findings of this study are available from the corresponding author upon reasonable request.

## Notes

### Competing Interest Statement

The authors have declared no competing interest.

