## Supporting Information for "Label-free Machine Learning Prediction of Chemotherapy on Tumor Spheroids using a Microfluidics Droplet Platform"

A. Jan

UMS3750 CNRS, Plateforme technologique Institut Pierre Gilles de Gennes, PSL Research University, 6 rue Jean Calvin, 75 005 Paris, France

S. Zaidi

INSERM U1330, Childrens' Oncology Research Unit, Research Center, PSL Research University, Institut Curie, 26 rue d'Ulm, 75 005 Paris, France

O. Delattre

SIREDO Oncology Center, Institut Curie, 26 rue d'Ulm, 75 005 Paris, France

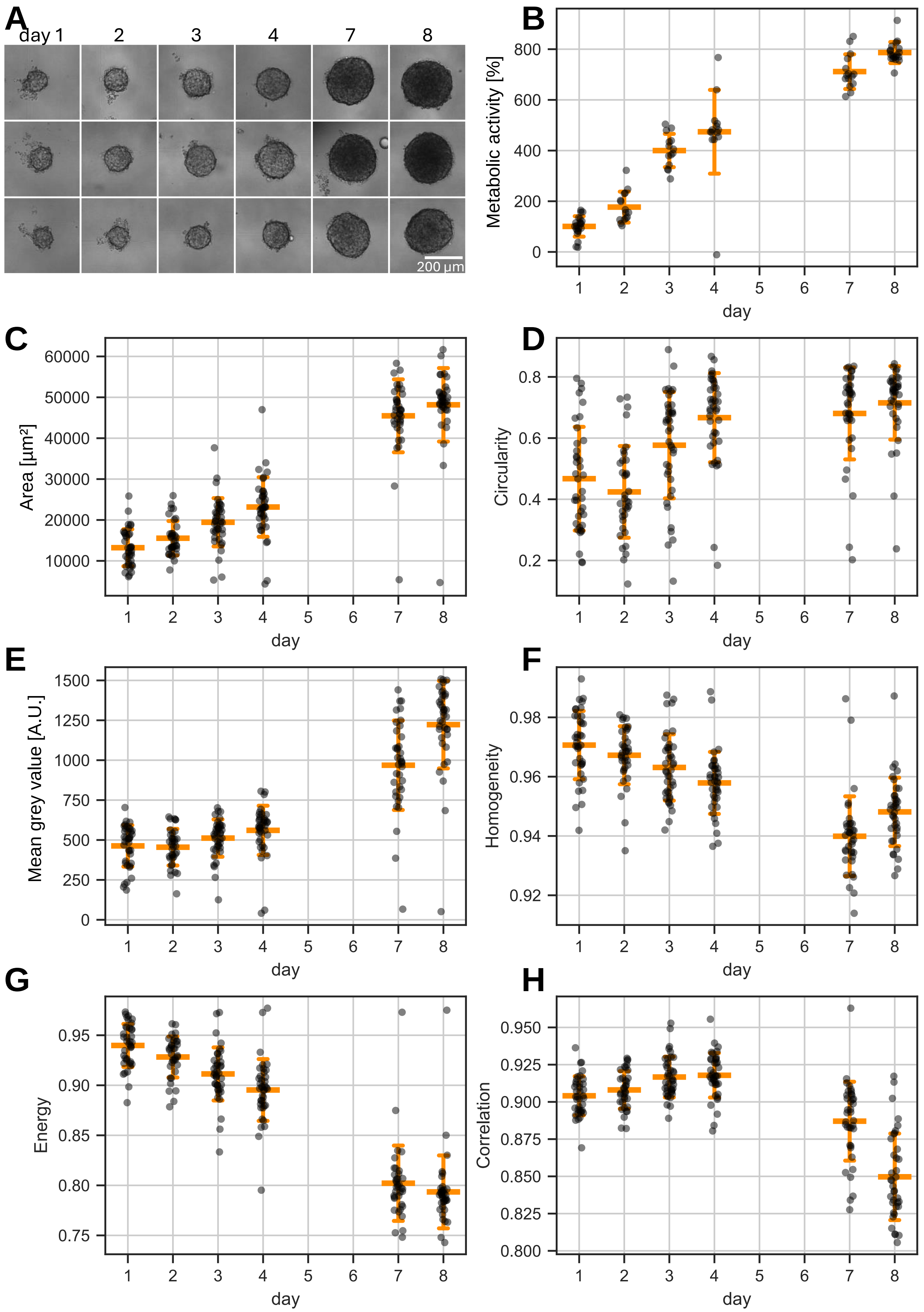


**Figure S1**. A) Images of PDX-derived spheroids over time. B) Normalized metabolic activity of spheroids over time. C-H. Morphological features of spheroids over time.

Spheroids were grown from 1000 cells in droplet. Each day, two tubes were imaged and one of them was used to perform metabolic activity assay using Alamar Blue. The horizontal bar represents the mean, the error bar represents the standard deviation of the mean.

| Feature [unit] | Description | Formula |
|---|---|---|
| Area [µm²] | Area of the spheroid cross-section. | |
| Perimeter [µm] | Perimeter of the spheroid cross-section. | |
| Equivalent diameter [µm] | Diameter of a perfect circle that would have the same area as the spheroid cross-section. | $\sqrt{\frac{4 \times area}{\pi}}$ |
| Circularity | Measure of how closely the spheroid cross-section resembles a perfect circle. | $\frac{4 \times \pi \times area}{perimeter^2}$ |
| Solidity | Measure of the compactness of the spheroid cross-section. | $\frac{area}{area\ convex\ Hull}$ |
| Mean grey value [A.U.] | Average intensity value of all the pixels within the spheroid cross-section, minus background. | |
| Homogeneity | Measure of how closely the gray levels of neighboring pixels are distributed in the image. Derived from GLCM matrix. [a)] | $\sum_{i,j=0}^{N-1} \frac{P_{i,j}}{1+(i-j)^2}$ |
| Energy | Measure of the uniformity or repetition of patterns in the spheroid. Derived from GLCM matrix. | $\sqrt{\sum_{i,j=0}^{N-1} P_{i,j}^2}$ |
| Correlation | Measure of the linear dependency between the gray levels of neighboring pixels. Derived from GLCM matrix. [b)] | $\sum_{i,j=0}^{N-1} P_{i,j} \left[ \frac{(i-\mu_i)(j-\mu_j)}{\sqrt{(\sigma_i^2)(\sigma_j^2)}} \right]$ |
| Growth [µm²] | Difference of area between two time point. | $area_{t2} - area_{t1}$ |
| Grey intensity variation [A.U.] | Difference of the mean grey value between two time point. | $grey\ value_{t2} - grey\ value_{t1}$ |

[a)] $P_{i,j}$ is the normalized GLCM value at position $(i, j)$. $N$ is the number of gray level in the image (8 bits so N = 256) ; [b)] $\mu_i = \sum_{i,j=0}^{N-1} i \times P_{i,j}$ , $\sigma_i = \sqrt{\sum_{i,j=0}^{N-1} P_{i,j} \times (i-\mu_i)^2}$

**Table S1**. Features extracted from spheroids images and their formula. The variables without formula were measured directly from the image.

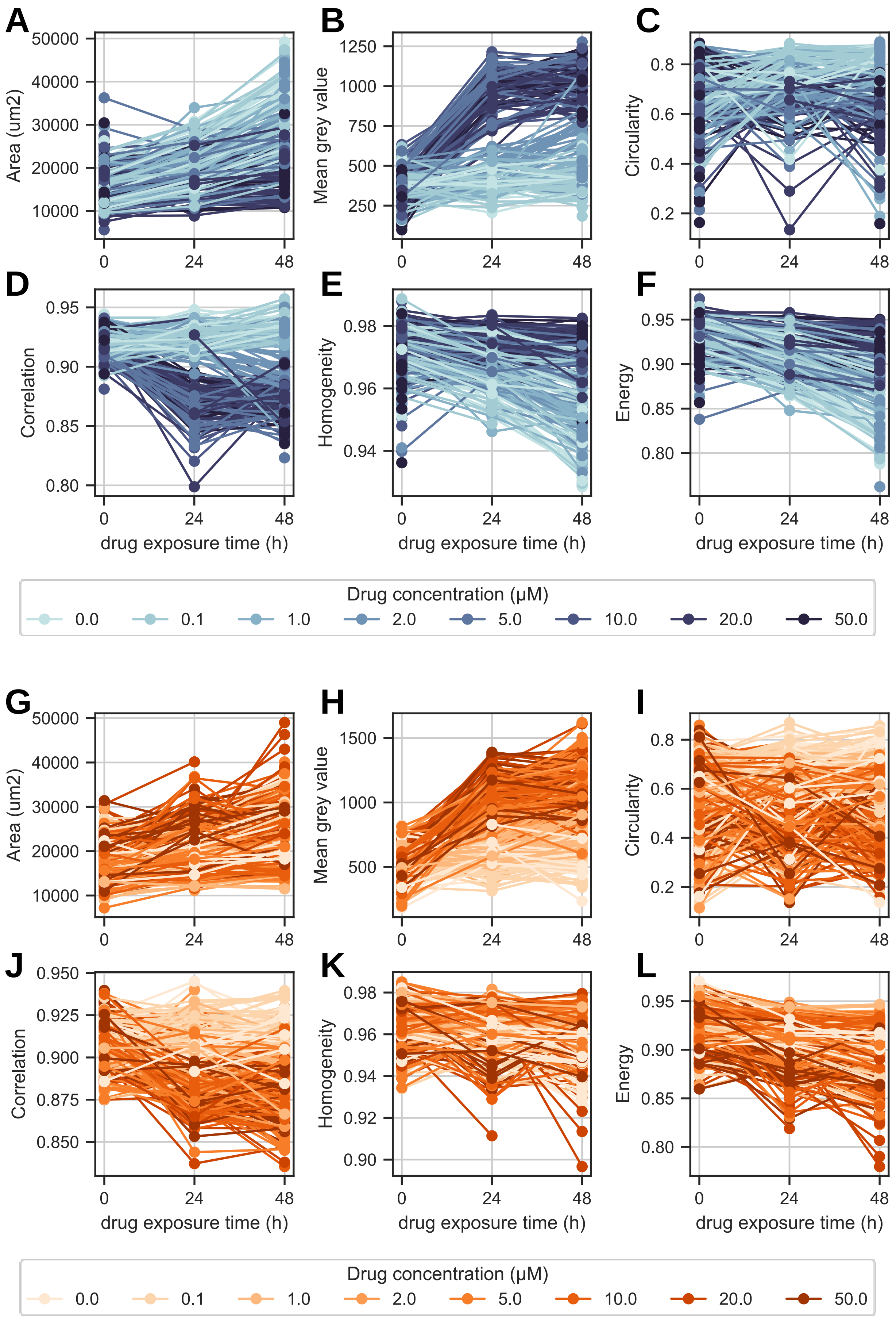


**Figure S2**. Morphological features of spheroids derived from A-F) cell line and G-L )PDX ; depending on time and drug concentration.

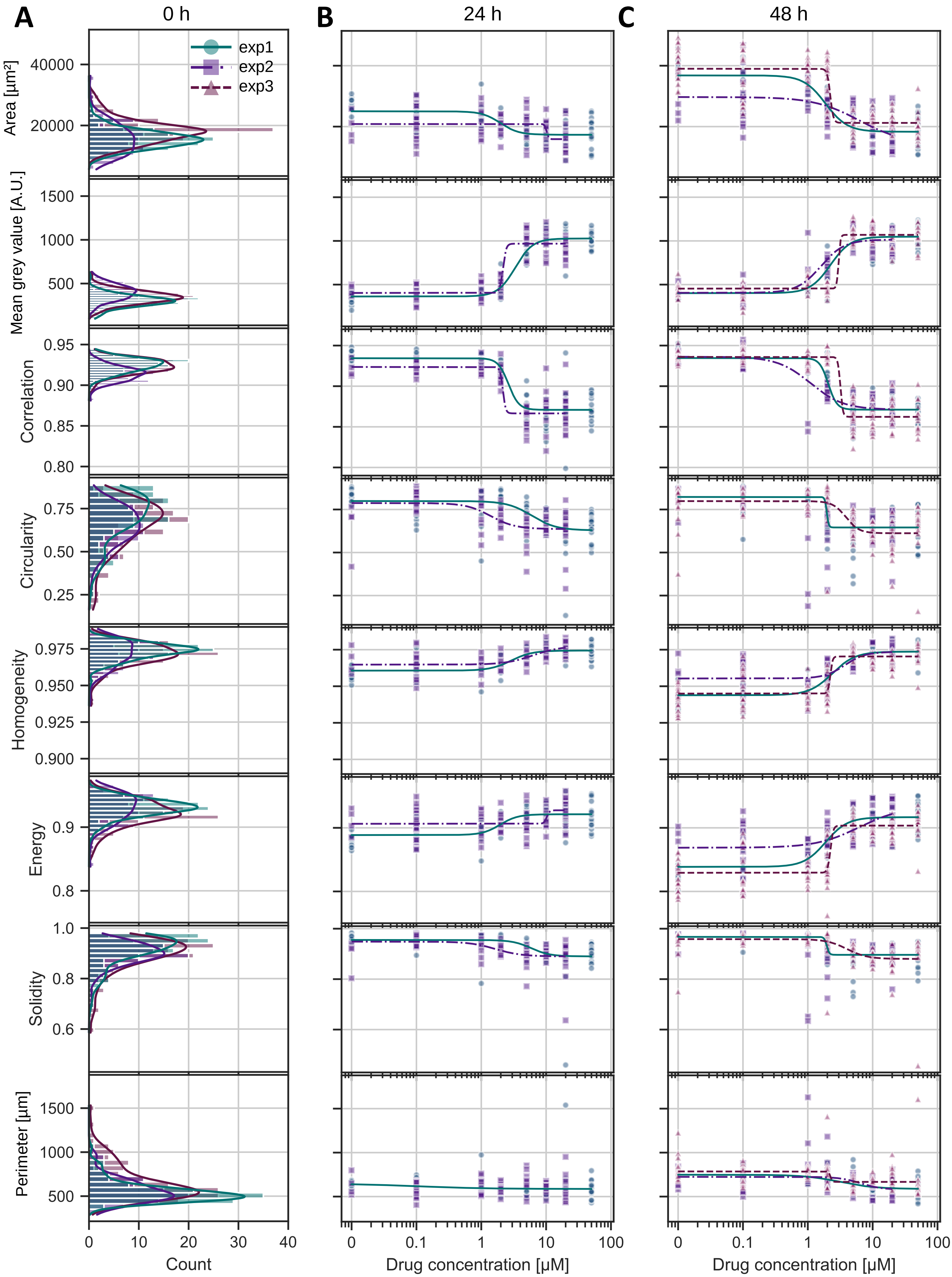


**Figure S3.** A) Histograms of morphological features of cell line-derived spheroids before drug exposure (0h). B & C) Morphological features of cell line-derived spheroids submitted to different drug concentrations after respectively 24 h and 48 h of drug exposure. Each point represents a single spheroid. Colors and shapes indicate independent experiments, with each unique combination corresponding to a different experiment. The curves represent sigmoid fits.

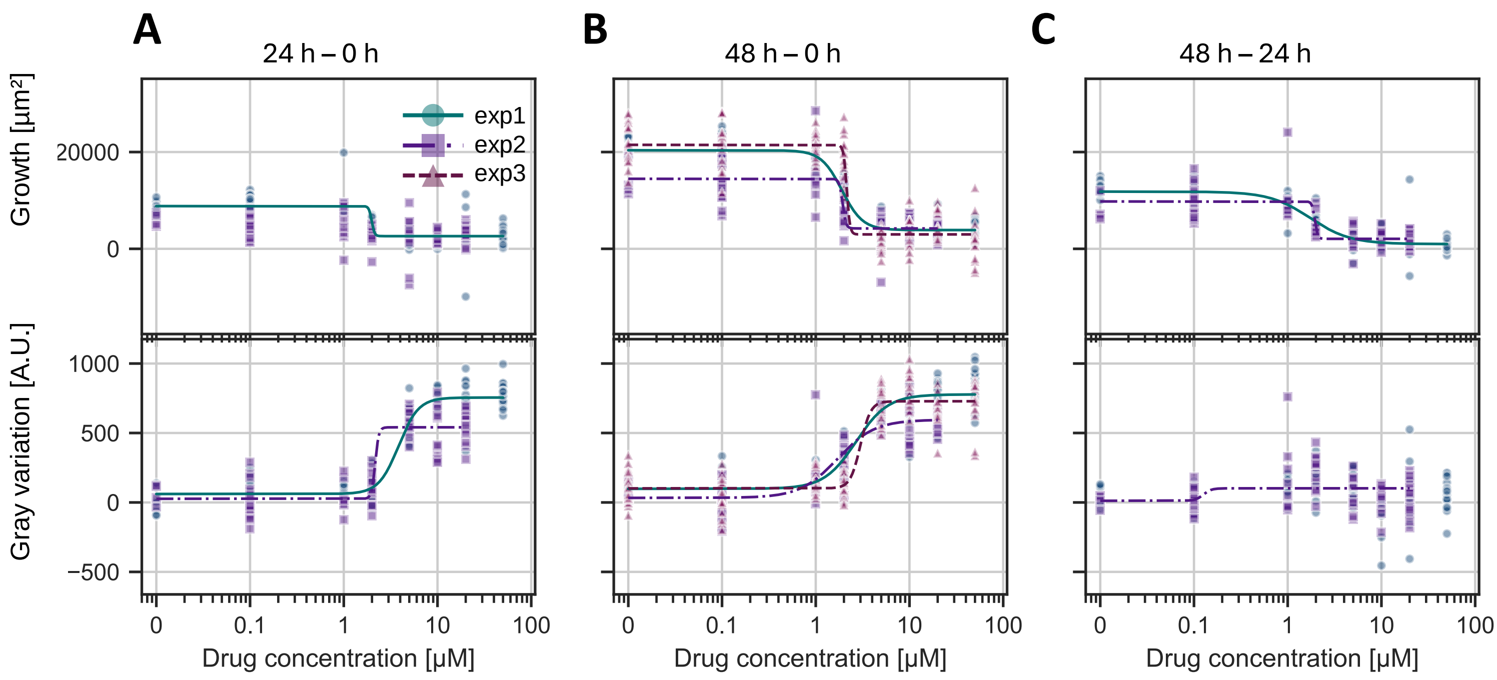


**Figure S4**. Variation of morphological features of cell line-derived spheroids submitted to different drug concentrations between A) 0 and 24 h ; B) 0 and 48 h and C) 24 and 48 h.

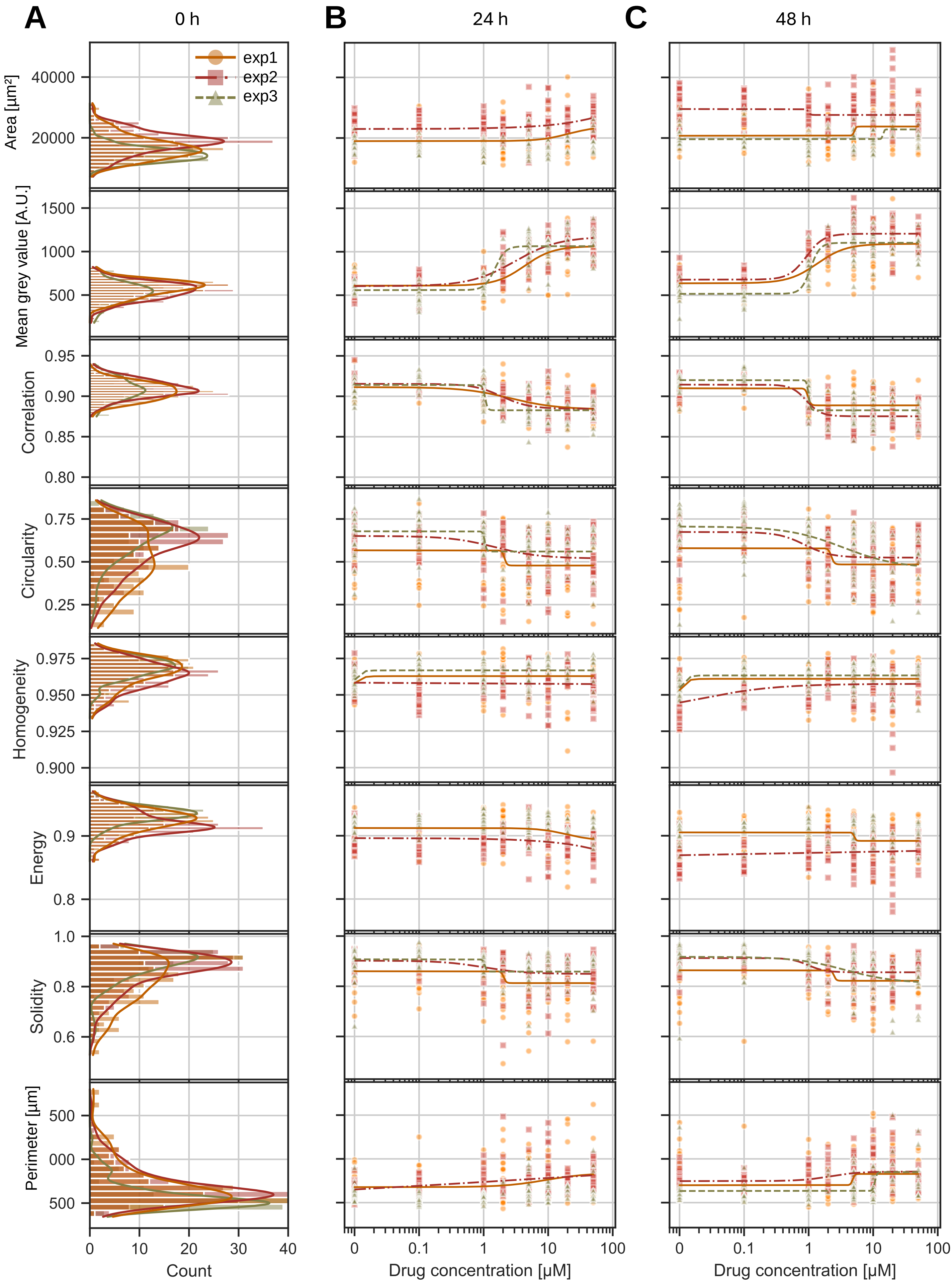


**Figure S5.** A) Histograms of morphological features of PDX-derived spheroids before drug exposure (0h). B) and C) Morphological features of PDX-derived spheroids submitted to different drug concentrations after respectively 24 h and 48 h of drug exposure. Each point represents a single spheroid. Colors and shapes indicate independent experiments, with each unique combination corresponding to a different experiment. The curves represent sigmoid fits.

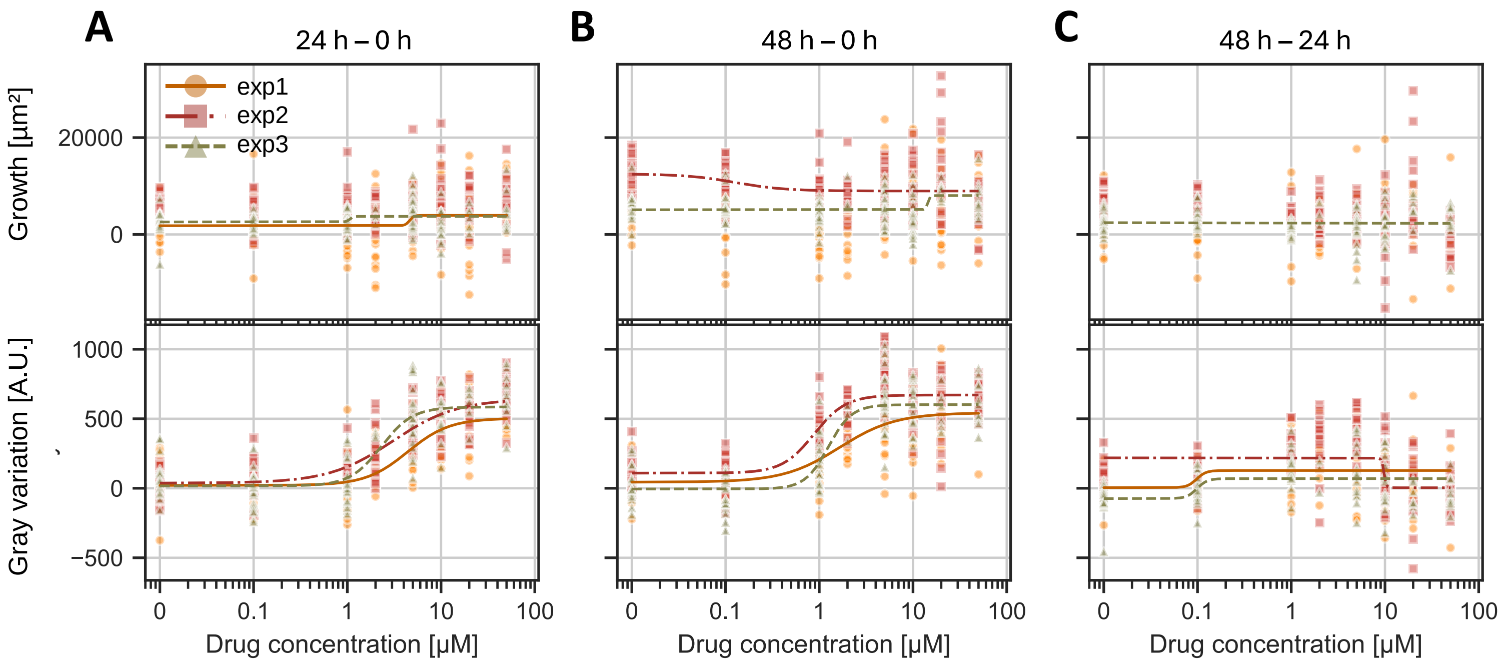


**Figure S6.** Variation of morphological features of PDX-derived spheroids submitted to different drug concentrations between A) 0 and 24 h ; B) 0 and 48 h and C) 24 and 48 h.

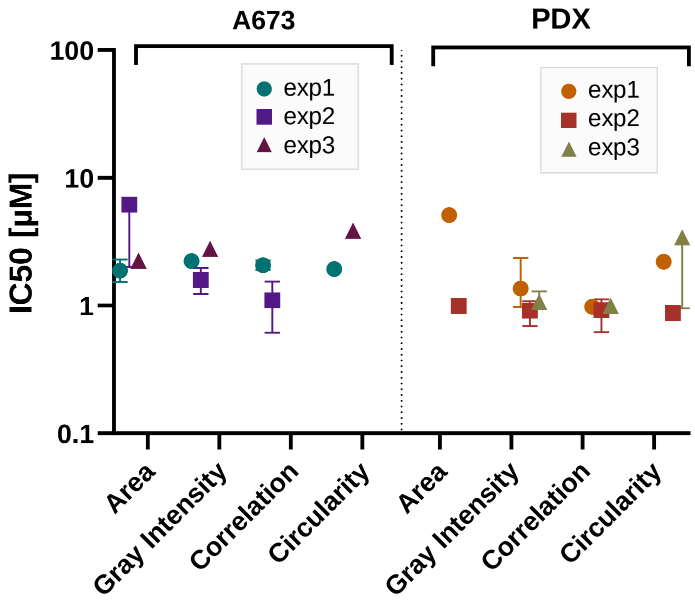


**Figure S7.** IC50 values derived from a sigmoid fit from feature VS drug curves. Error bar represents the 95% CI of the IC50 value. No error bar means poor fit.

Fig S8

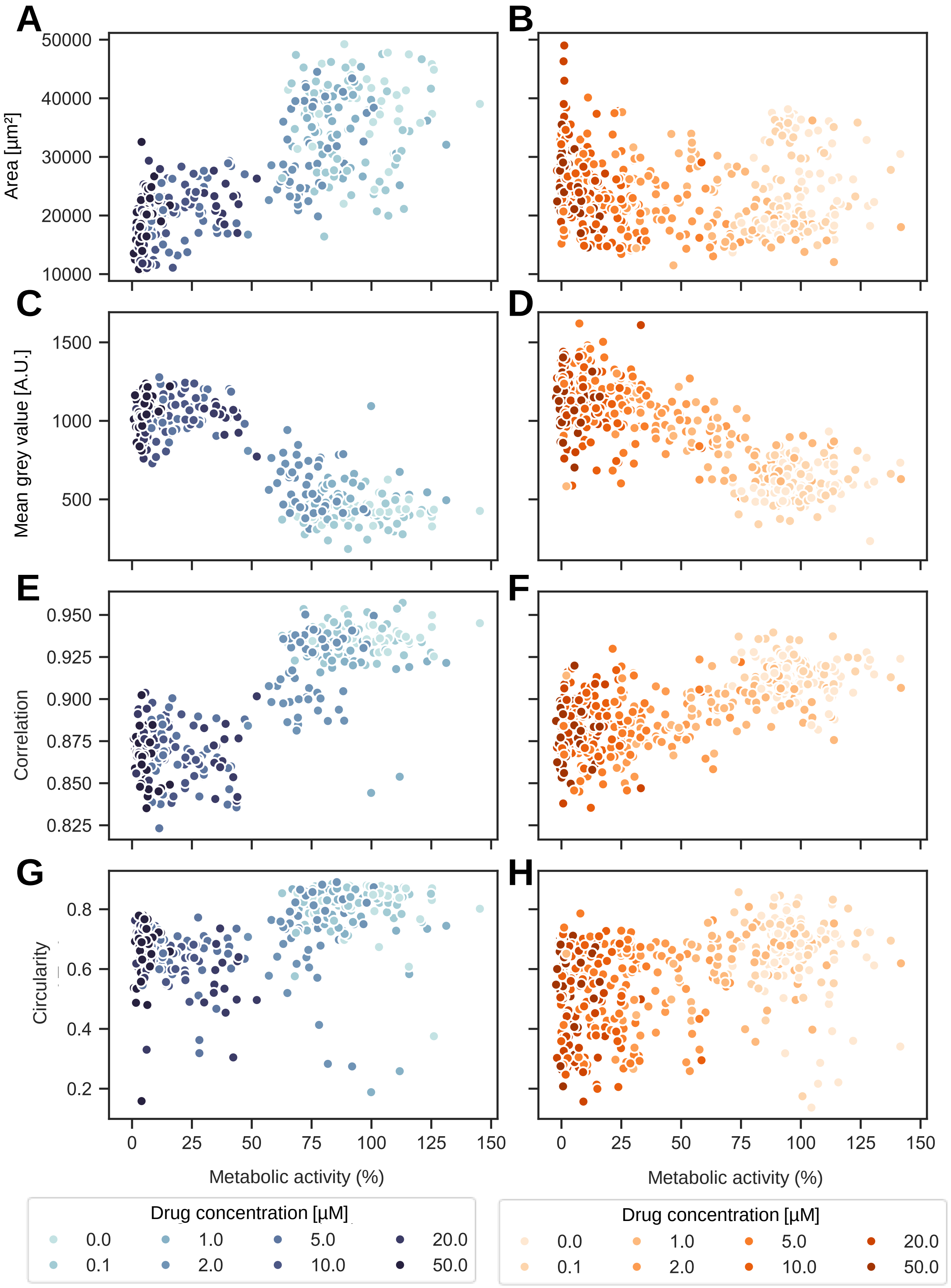


**Figure S8.** Features depending on the metabolic activity for A, C, E, G) cell line and B, D, F, H) PDX after 48 h of drug treatment. Each point represents one spheroid, its color represents the drug concentration.

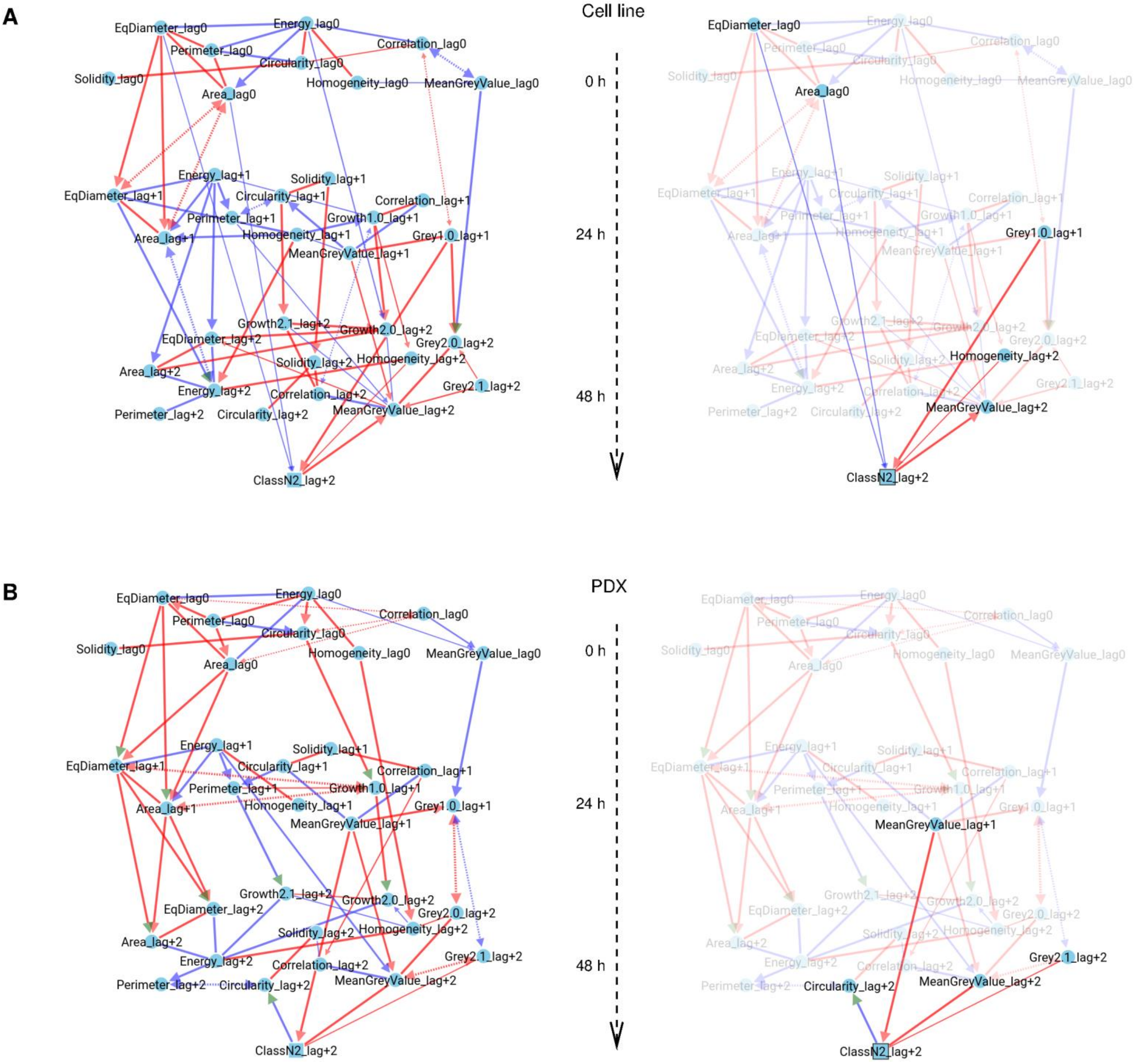


**Figure S9.** MIIC network for A) cell line and b) PDX. Left: full network with temporal orientation. Right: direct links with target class highlighted.

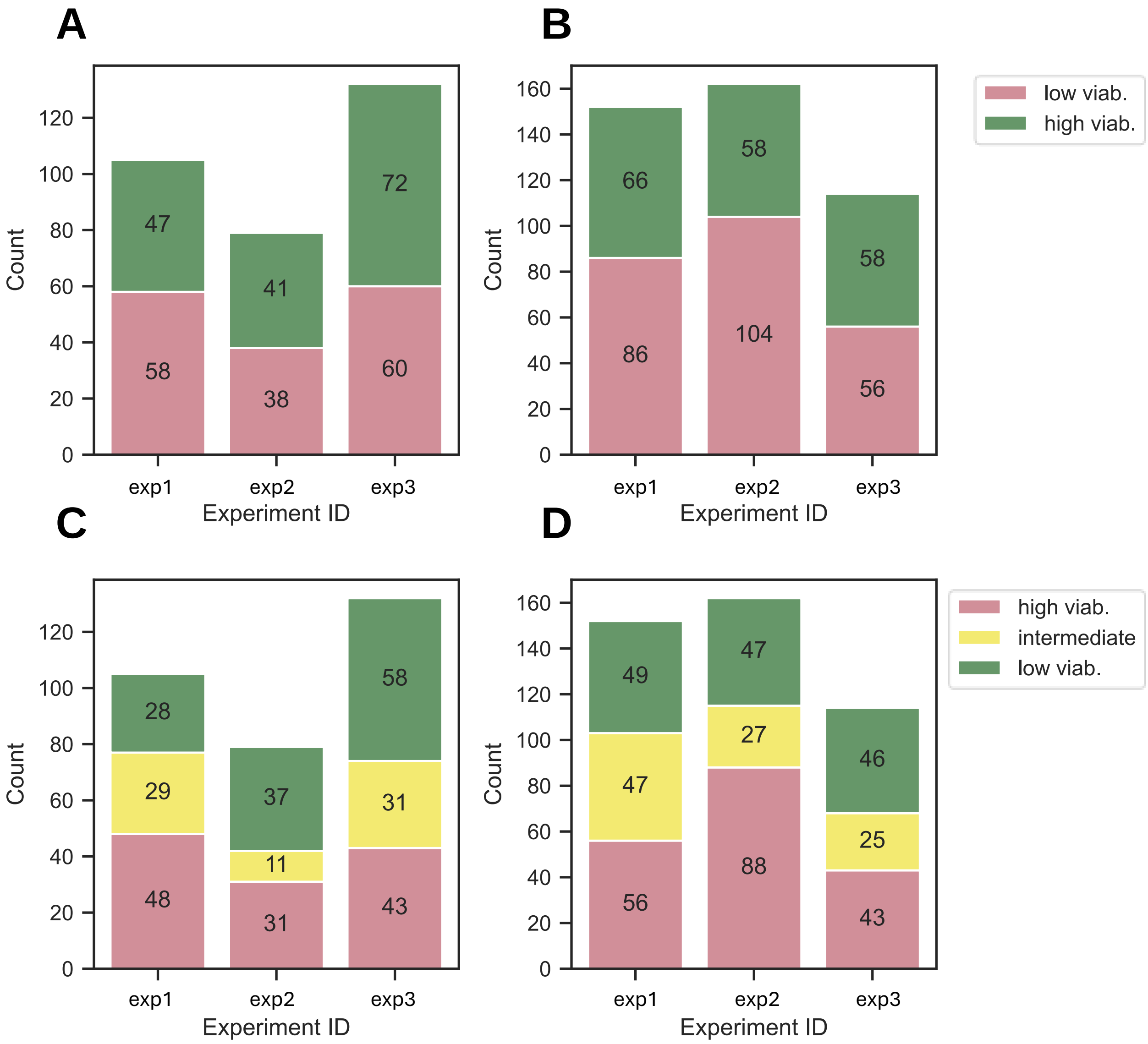


**Figure S10.** Description of the dataset compositions used for machine learning. A) cell line dataset, 2 classes. B) PDX dataset, 2 classes. C) cell line dataset, 3 classes. D) PDX dataset, 3 classes.

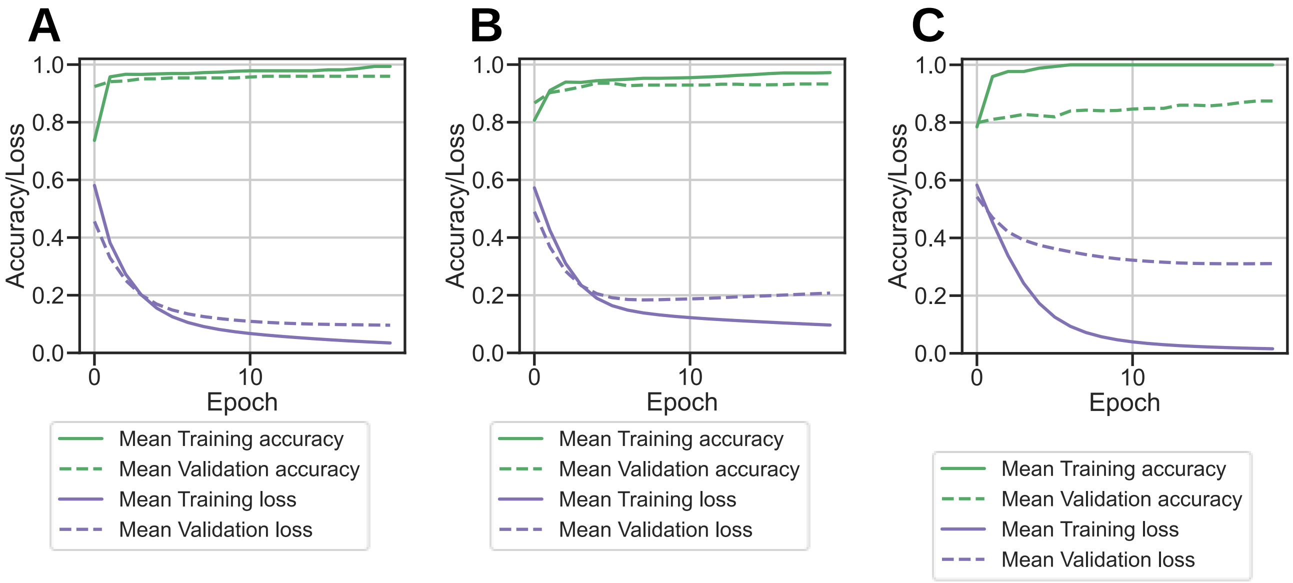


**Figure S11.** Accuracy/Loss vs. epoch. A) Train and Test on cell line dataset, B) Train and Test on PDX dataset & C) Train on cell line dataset, Test on PDX dataset. Epoch = 20, batch size = 30.

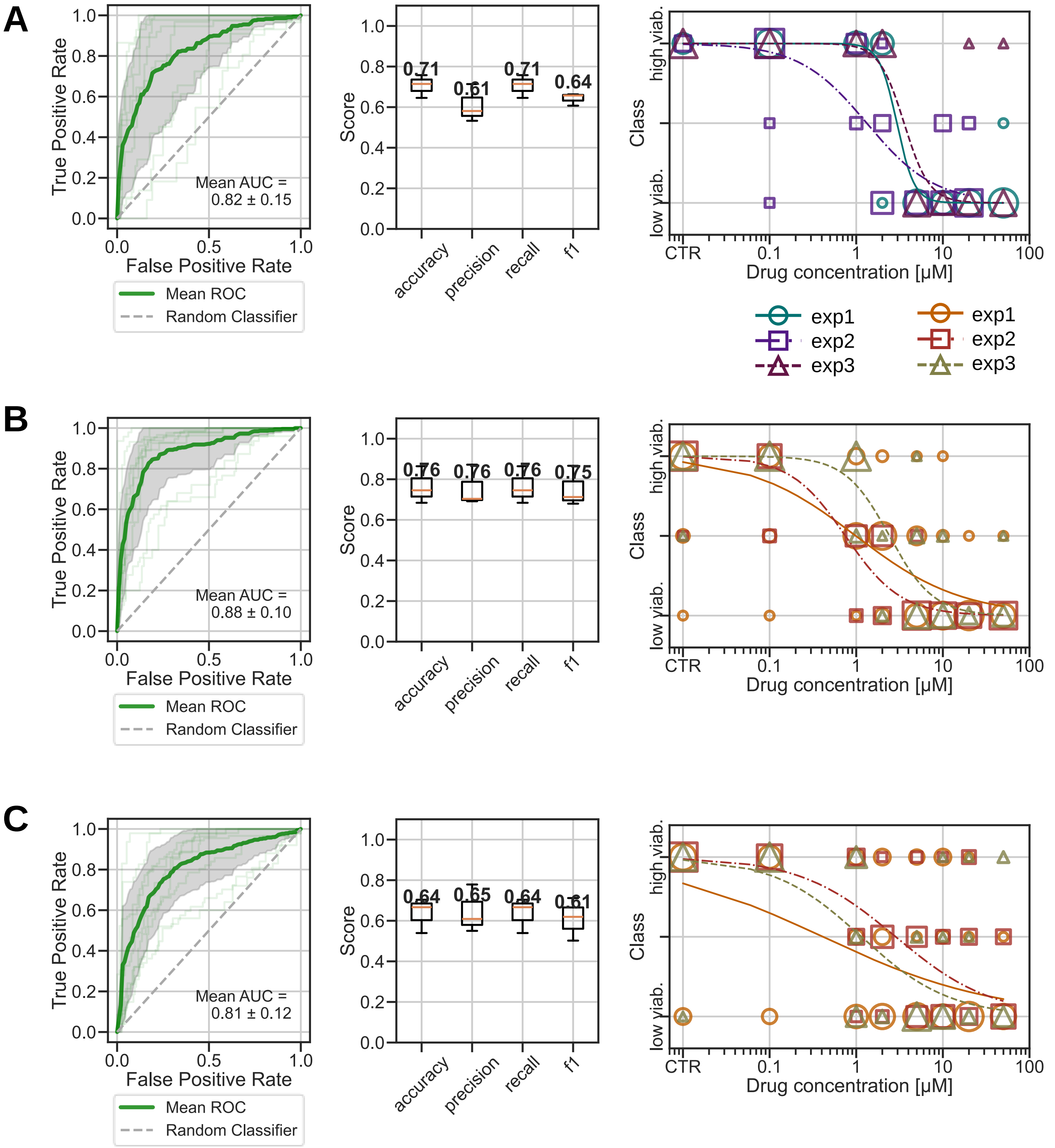


**Figure S12.** Results of machine learning model for 3 classes classification. Train/test on A) cell line, B) PDX dataset. C) Train on cell line, test on PDX dataset. For each, ROC-AUC curve, metrics and dose-response curves predictions are shown.

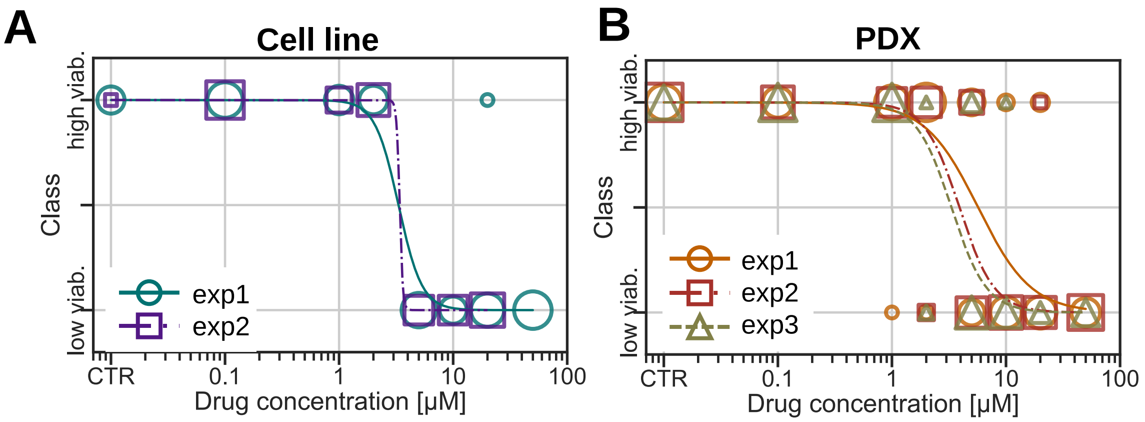


**Figure S13.** Predicted dose-response curves after 24 h of treatment for A) cell line and B) PDX spheroids in droplets, derived from predictions made by the neural network machine learning model. The model was trained on morphological parameters measured after 0 and 48 hours of drug exposure and predicted the viability classes at 24 hours. The curves represent sigmoid fits. Each combination of color and shape represent one independent experiment. The size of each point on the plot is scaled proportionally to the number of overlapping data points.

| Cell type | Experiment | IC50 48h [µM] 95% CI [a] | IC50 24h [µM] 95% CI |
|---|---|---|---|
| Cell line | 1 | 3.4 [2.3, 4.7] | 3.3 [2.5, 4.3] |
| | 2 | 2.4 [2.1, 3.4] | 3.4 [2.3, 4.6] |
| | 3 | 3.4 [2.8, 4.2] | x |
| PDX | 1 | 2.1 [1.7, 2.6] | 5.7 [4.3, 7.5] |
| | 2 | 1.1 [1.0, 15.0] | 3.9 [3.1, 4.8] |
| | 3 | 1.8 [1.4, 2.5] | 3.3 [2.4, 4.4] |

[a] Values from dose-response curves issued from viability classes after metabolic activity discretization

**Table S2**. Comparison of IC50 values obtained from the fit of the viability class after 48 h of treatment with the predicted viability classes after 24 h of treatment.

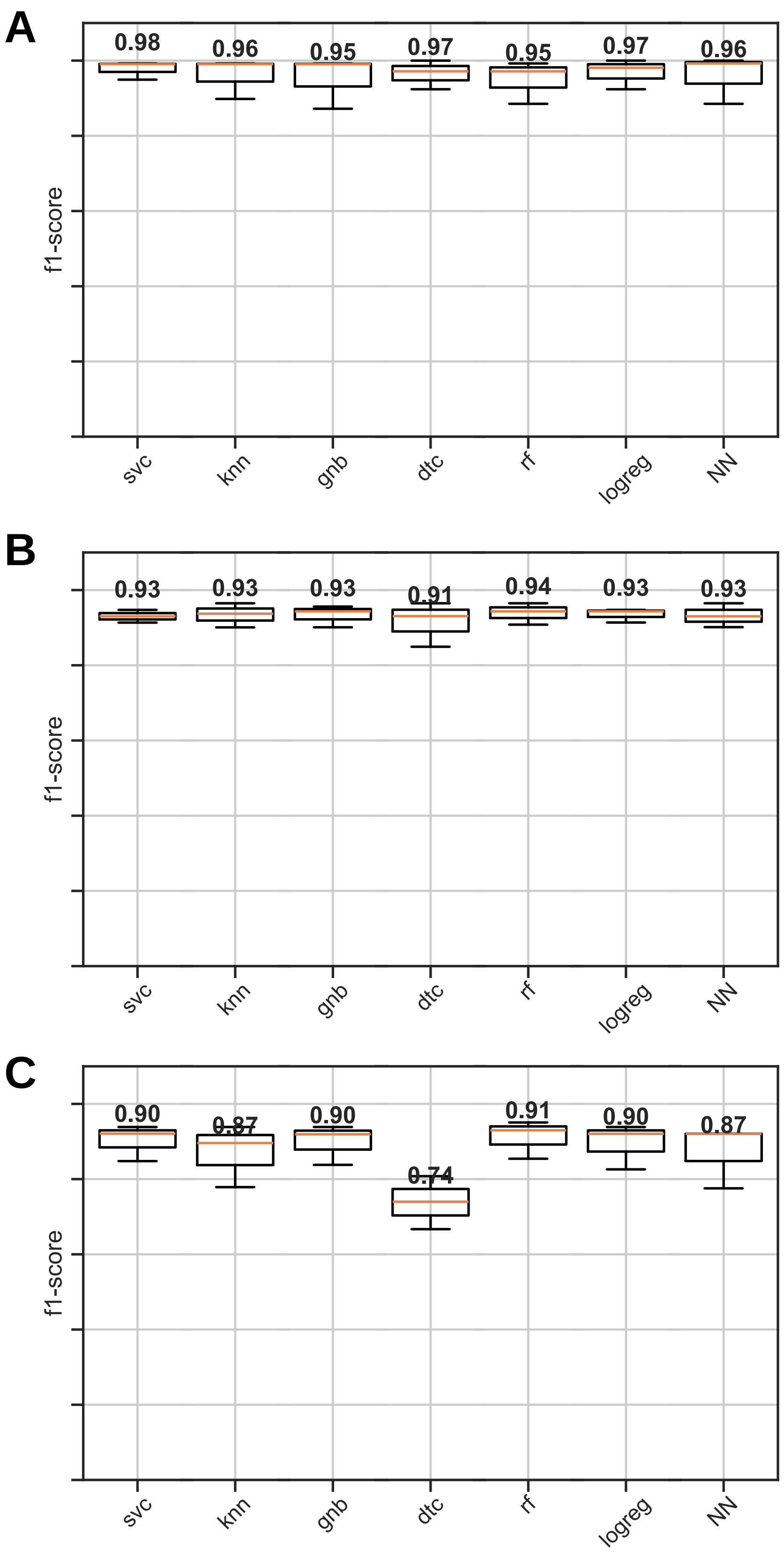


**Figure S14.** Comparison of f1-scores obtained with other machine learning models for the classification into 2 classes for A) cell line, B) PDX and C) training on cell line and test on PDX.
svc: Support Vector Classification ; knn: k-Neighbors ; gnb: Gaussian Naive Bayes ; dtc: Decision Free Classifier, rf: Random Forest, logreg: Logistic Regression ; NN: Neural Network.
The parameters used for each models are presented Table S3.

| Model | | A) Train / Test Cell line | B) Train / Test PDX | C)Train cell line / Test PDX |
|---|---|---|---|---|
| svc | C | 0.1 | 0.1 | 0.1 |
| | kernel | linear | rbf | linear |
| | gamma | 10 | 0.01 | 10 |
| knn | n_neighbors | 5 | 13 | 3 |
| | weights | uniform | uniform | uniform |
| | metric | manhattan | manhattan | manhattan |
| gnb | var_smoothing | 1e-9 | 1e-9 | 1e-9 |
| dtc | criterion | gini | entropy | gini |
| | max_depth | None | 30 | None |
| | min_samples_split | 10 | 10 | 2 |
| | min_samples_leaf | 2 | 5 | 1 |
| rf | n_estimators | 50 | 50 | 100 |
| | max_depth | 6 | 6 | 6 |
| | min_samples_split | 2 | 10 | 10 |
| | min_samples_leaf | 1 | 5 | 5 |
| | max_features | sqrt | log2 | log2 |
| logreg | penalty | l2 | l2 | l2 |
| | C | 100 | 0.01 | 0.1 |
| | max_iter | 100 | 100 | 100 |

**Table S3.** Models hyperparameters used to test each model and each dataset.
Each model was optimized, using GridSearchCV to select the best hyperparameters explored using predefined grids. As a cross-validation, models were trained on two out of three experiments and tested on the remaining one.

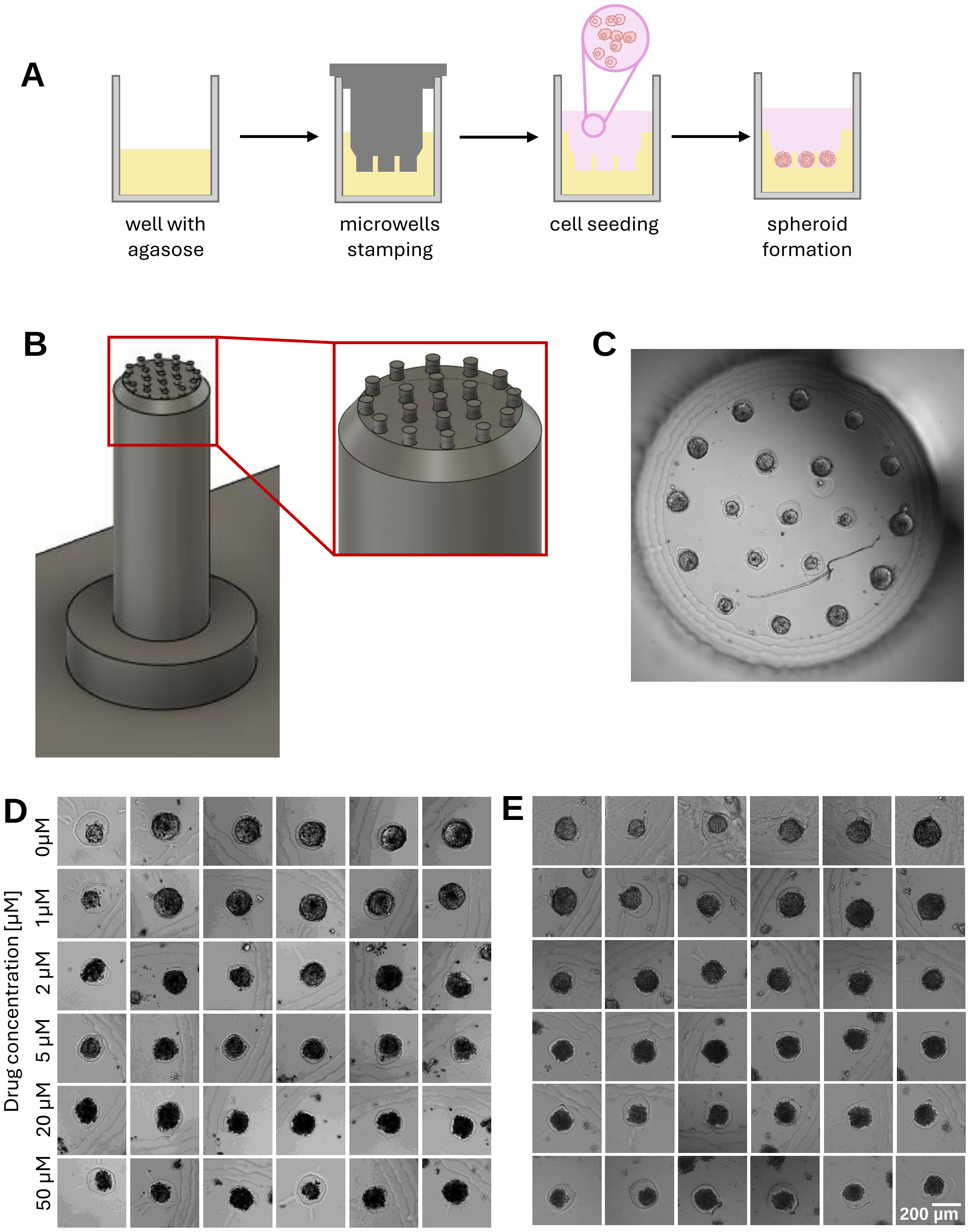


**Figure S15.** A) Protocol to produce spheroids in agarose microwells.
B) 3D-model of the mold used to produce the microwells.
C) Image of a well containing 19 spheroids, each in a microwell.
D) & E) Images of spheroids in microwells derived from cell line and PDX respectively, after 48 h of drug (Etoposide) treatment.

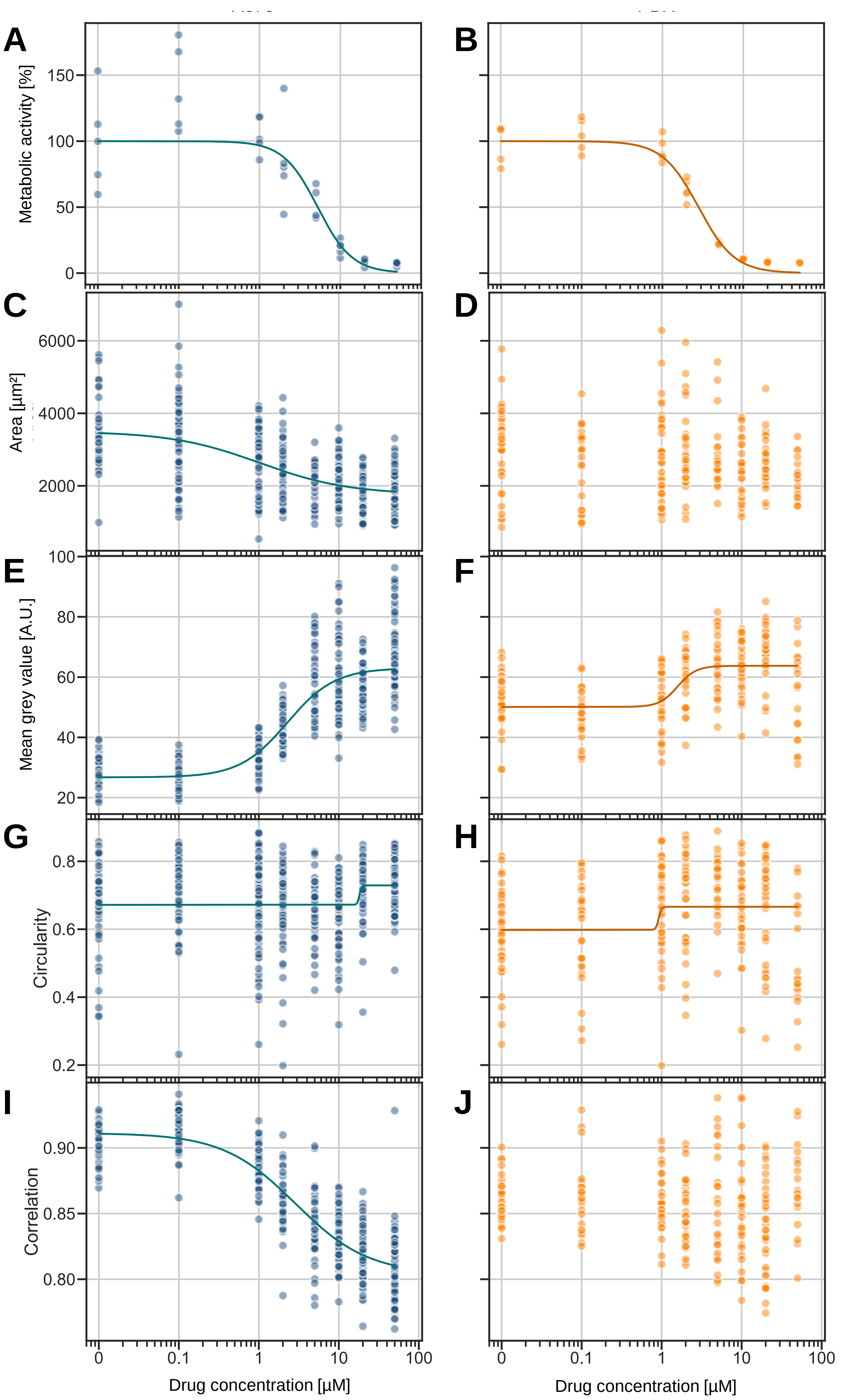


**Figure S16.** A, B) Metabolic activity of cell line and PDX spheroids in microwells respectively, after 48 h of drug treatment. C, E, G, I) and D, F, H, J) Morphological features of respectively cell line and PDX-derived spheroids in microwells after 48 h of drug treatment. Each point represent a spheroid, the line is a sigmoid fit to the points.

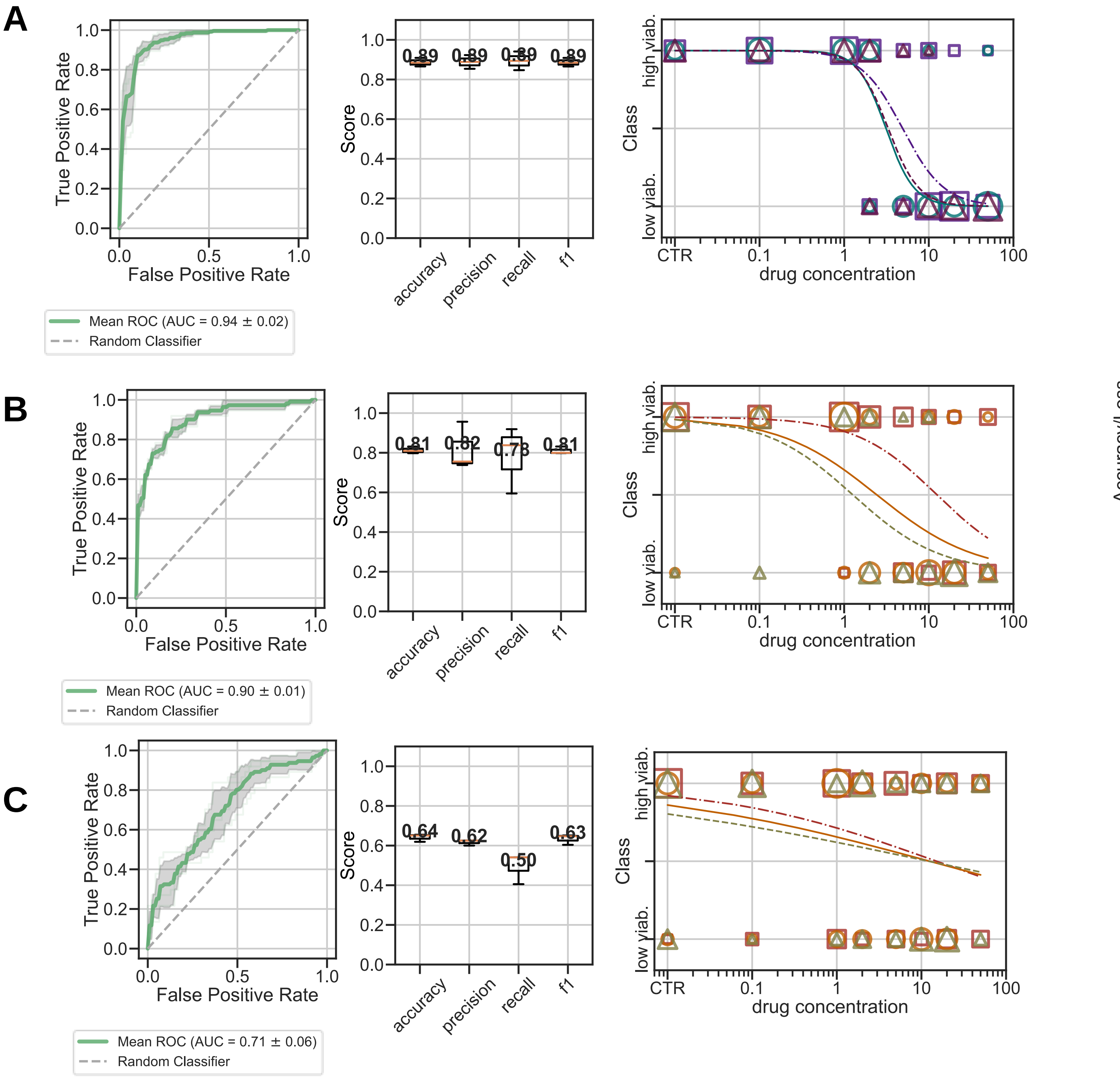


**Figure S17.** Results of machine learning model classification for spheroids grown in microwells. Train/test on A) cell line, B) PDX dataset. C) Train on cell line, test on PDX dataset. For each, ROC-AUC curve, metrics and dose-response curves predictions are shown.

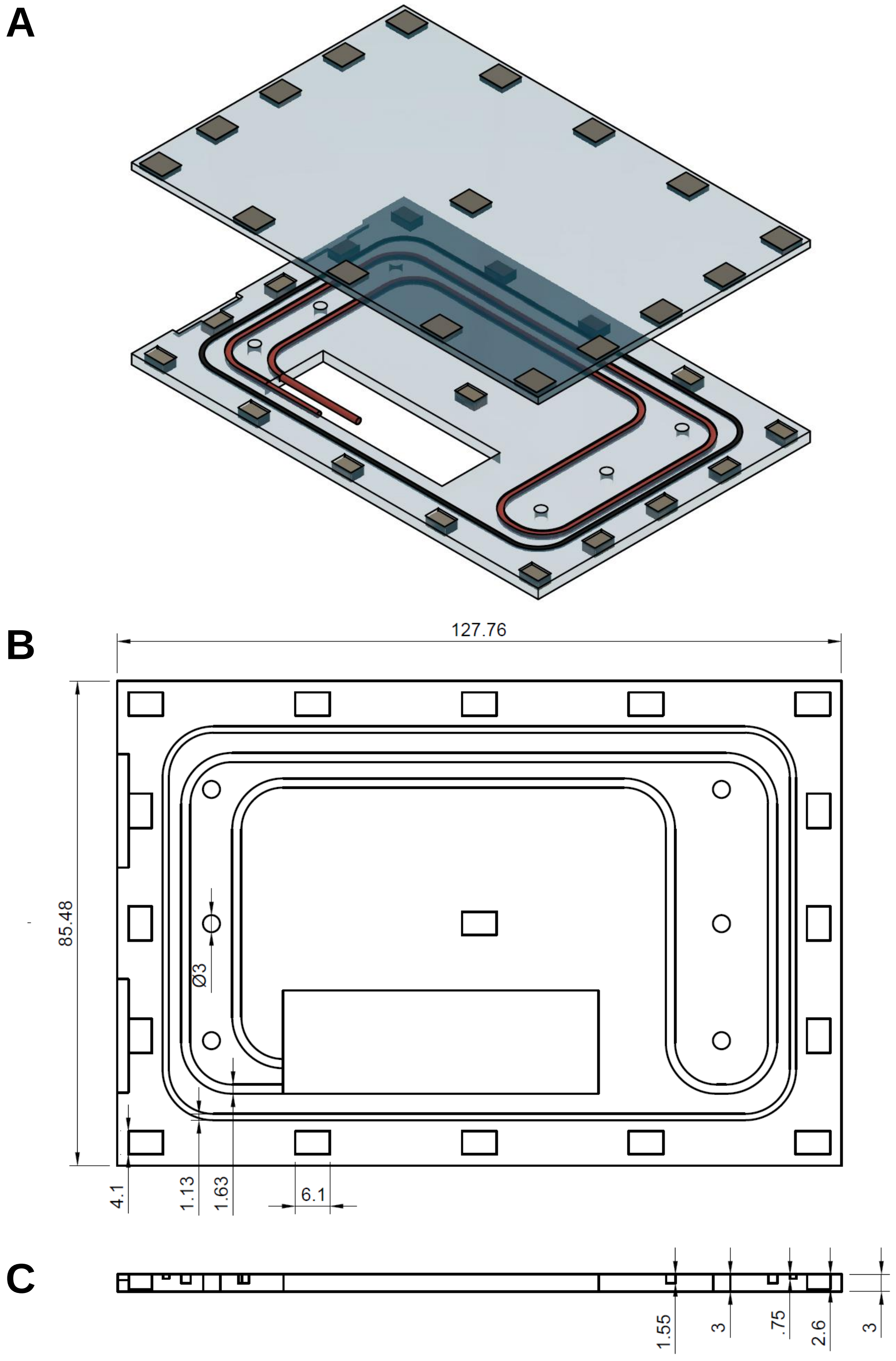


**Figure S18.** Schematic of the cartridge for spheroids imaging in tubing . A) Exploded view showing the two components, on top: plate to image through (thickness 2mm), bottom: plate to hold tubing (red), with a rubber joint (black). Each plate contains 17 embedded magnets for secure assembly. B) and C) Front and lateral view respectively of the tube holder plate, with dimensions in millimeters.

### Supplementary Methods.

#### Spheroid production & drug screening in microwells

A 2% agarose gel in PBS was prepared by dissolving agarose (Sigma-Aldrich, A0576) in PBS and heating the solution. Then, 200 µL of liquid agarose was pipetted into each well of a 96-well plate (Revvity, #6055302). While the agarose was still liquid, a 3D-printed mold was applied to create the microwells. Once the agarose had jellified, the mold was removed, and the plate was exposed to UV light for disinfection. The 3D-printed mold consisted of 19 cylindrical pillars, each with a diameter of 200 µm and a height of 200 µm, evenly distributed within a single well of a 96-well plate.

Then, 25 µL of cells (600 000 cells $ml^{-1}$ for the cell line and 1 600 000 cells $mL^{-1}$ for the PDX) were added into each well. After waiting for the cells to sediment, 100 µL of the appropriate cell culture media was added, and the plate was incubated at 37°C, under 5% $CO_2$ in humidified atmosphere. After 24 hours, 25 µL of the media was removed and 30 µL of etoposide solution was added in the desired concentrations. After 48 hours of drug exposure, 13 µL of alamarBlue was added to each well.

The wells were imaged using plate reader with 1600 µm global focus height and focus offset of 20 µm with 6% excitation power and 6ms of exposure time. Images were taken after 48 h of drug exposure.

#### Image analysis of spheroids in microwells

Each spheroid was identified manually. Then, a program automatically selected the best focal plan and segmented the spheroid to extract its morphological properties, as for the droplet's experiments. The features that were extracted from the images were area, circularity, mean gray level, homogeneity, energy and correlation.

#### Metabolic assay in microwells

After overnight incubation with Alamar Blue reagent, the fluorescent signal of each well was measured using the plate reader with excitation wavelength of 560nm, emission wavelength of 590 nm, measurement height of 9.5nm and 100 flashes. The values were normalized by the control and the background subtracted, as for droplet experiment.

#### Label-free analysis of spheroids in microwells

The same neural network model as the one used for the droplet experiments was applied to the microwells dataset. The features used were the area, circularity, mean grey level, and textural parameters (energy, correlation, homogeneity).

Only one experiment was performed per cell type. The dataset was composed of n = 494 for the cell line, and n = 251 for the PDX.

Each dataset was split randomly into a training and a testing set using a 3-fold cross-validation process. For each dataset (cell line and PDX, the model was then trained on 2 out of the 3 split and tested on the third split. The same metrics as for the droplets were measured to evaluate the accuracy of the classification.

The model was then trained on cell line dataset and tested on PDX dataset, split randomly in 3 parts.
